# Phosphatidylinositol 5 phosphate 4-kinase regulates plasma-membrane PIP_3_ turnover and insulin sensitivity in *Drosophila*

**DOI:** 10.1101/333153

**Authors:** Sanjeev Sharma, Swarna Mathre, Sanjeev Visvanathan, Dhananjay Shinde, Padinjat Raghu

## Abstract

Phosphatidylinositol-3,4,5-trisphosphate (PIP_3_) generation at the plasma membrane is a key event during activation of receptor tyrosine kinases such as the insulin receptor and is critical for normal growth and metabolism. The lipid kinases and phosphatases regulating PIP_3_ levels are described but mechanisms controlling their activity remain unclear. We report that in *Drosophila*, phosphatidylinositol 5 phosphate 4-kinase (PIP_4_K) function at the plasma membrane is required for normal PIP_3_ levels during insulin receptor activation. Depletion of PIP_4_K increases PIP_3_ levels and augments sensitivity to insulin through enhanced Class I phosphoinositide 3-kinase (PI_3_K) activity. Animals lacking PIP_4_K show enhanced insulin signalling dependent phenotypes *in vivo* and are resistant to the metabolic consequences of a high-sugar diet, highlighting the importance of PIP_4_K in normal metabolism and development. Thus, PIP_4_KS are key regulators of receptor tyrosine kinase signalling with implications for growth factor dependent processes including tumour growth, T-cell activation and metabolism.

## Introduction

Lipid kinases that can phosphorylate selected positions on the inositol head group of phosphatidylinositol (PI), generate second messengers that regulate multiple processes in eukaryotic cells. The generation of phosphatidylinositol 3,4,5-trisphosphate (PIP_3_) through the action of Class I PI_3_K following growth factor receptor (e.g Insulin receptor) stimulation, is a widespread signalling reaction [1] that regulates normal growth and development [2]. The role of Class I PI_3_K activation in response to insulin receptor signalling is evolutionarily conserved and has been widely studied in metazoan models such as the fly, worm and mammals [3]. Robust control of the levels and the dynamics of PIP_3_ turnover is essential to maintain fidelity and sensitivity of information transfer during insulin signalling. This is achieved through a number of different molecular mechanisms. The Class I PI_3_K enzyme is a dimer of a catalytic subunit (p110) whose activity is inhibited under unstimulated conditions by the regulatory subunit (P85/50/55/60). Upstream receptor activation and subsequent binding to p-Tyr residues on the receptor and adaptor proteins relieves this inhibition. In addition, lipid phosphatases are also important in controlling PIP_3_ levels at the plasma membrane. PTEN, a 3-phosphatase, hydrolyzes PIP_3_ to produce PI(4,5)P_2_ [4] while SHIP_2_ is a 5-phosphate that generates PI(3,4)P_2_ from PIP_3_ [5]. It is well documented that mutations in genes encoding any of these enzymes can be oncogenic or result in metabolic syndromes. Loss of function in PTEN or gain of function in Class I PI_3_K genes results in tumour development [6] while loss of SHIP2 results in altered insulin sensitivity in mammals [7,8]. Thus, the control of receptor-activated PIP_3_ levels is vital to the regulation of events that direct cell growth and metabolism.

Class I PI_3_K enzymes utilize phosphatidylinositol 4,5-bisphosphate [PI(4,5)P_2_] as substrate to generate PIP_3_. In animal cells, the major route of PI(4,5)P_2_ synthesis is the action of phosphatidylinositol 4 phosphate 5-kinase (PIP_5_K), enzymes that use phosphatidylinositol 4-phosphate (PI_4_P) as substrate and phosphorylate position 5 of the inositol headgroup [9]. More recently, Cantley and colleagues have described a distinct class of lipid kinases, the phosphatidylinositol 5 phosphate 4-kinases (PIP_4_K), enzymes that utilize phosphatidylinositol 5-phosphate (PI_5_P) as substrate and phosphorylate position 4 to generate PI(4,5)P_2_ [10]. Loss of PIP_4_Ks does not result in a drop in the mass of total cellular PI(4,5)P_2_ but the levels of its preferred substrate, PI_5_P are elevated [[11], reviewed in [12]]. In mammalian cells, three isoforms of PIP_4_KS occur, viz. PIP_4_K_2_A, PIP_4_K_2_B and PIP_4_K_2_C. The phenotypes of mouse knockouts in each of these genes suggest a role for PIP_4_KS in regulating receptor tyrosine kinase and PI_3_K signaling; deletion of PIP_4_K_2_A and PIP_4_K_2_B is able to slow tumor growth in ps3−/− mice [13]; depletion of PIP_4_K_2_C results in excessive T-cell activation [14] and loss of PIP_4_K_2_B in mice results in hyper-responsiveness to insulin and a progressive loss of body weight in adults [15]. Previous studies have linked PIP_4_K_2_B to insulin and PI_3_K signalling. Overexpression of PIP_4_K_2_B in CHO-IR cells (expressing extremely low levels of endogenous PIP_4_K_2_B) results in reduced PIP_3_ production following insulin stimulation [16]. Similarly, in U20S cells, acute doxycycline-induced overexpression of PIP_4_K_2_A reduces AKT activation seen on insulin stimulation although changes in PIP_3_ levels were not reported under these conditions [17]. By contrast, a recent study has reported that in immortalized B-cells that carry a deletion of PIP_4_K_2_A, there is a reduction in PIP_3_ levels following insulin stimulation [18]. Thus, although there are multiple lines of evidence suggesting a link between PIP_4_K and Class I PI_3_K signaling during insulin stimulation, the impact of the PIP_4_K function on PIP_3_ levels remains unresolved.

It has been reported that loss of the only PIP_4_K in *Drosophila* results in a larval growth deficit and developmental delay. These phenotypes were associated with an overall reduction in the levels of pS6K^T398^ and pAKT^S505^, both outputs of <u>*m*</u>echanistic <u>*T*</u>arget <u>*O*</u>f <u>*R*</u>apamycin (mTOR) signalling. The systemic growth defect in the dPIP_4_K mutants (*dPIP_4_K^29^*) could be rescued by enhancing mTOR complex 1(TORC_1_) activity through pan-larval overexpression of its activator Rheb [11,19]. Since then it has also been shown in mice that PIP_4_K_2_C can regulate TORC_1_-mediated signalling in immune cells [14]. The loss of PIP_4_K_2_C was also shown to enhance TORC_1_ outputs in Tsc1/2 deficient MEFs during starvation [20]. mTOR signalling can transduce multiple developmental and environmental cues including growth factor signalling, amino acid and cellular ATP levels into growth responses [21]. However, the relationship between PIP_4_K function and its role in regulating TORC_1_ activity and Class I PI_3_K signaling remains unclear.

During *Drosophila* development, larval stages are accompanied by a dramatic increase in body size. Much of this growth occurs without increases in cell number but via an increase in cellular biomass that occurs in polyploid larval tissues such as the salivary gland and fat body [22]. One major mechanism that drives this form of larval growth is the ongoing insulin signalling; characterized by the endocrine secretion of insulin-like peptides (dlLPs) from insulin-producing cells (IPCs) in the larval brain and their action on peripheral tissues through the single insulin receptor in flies [23]. Removal of insulin receptor (*dInR*) activity [24] or the insulin receptor substrate (*chico*) [25] results in reduced growth and delayed development through multiple mechanisms. In flies, cell size in the salivary glands can be tuned by enhancing cell-specific Class I PI_3_K-dependent PIP_3_ production [26]. In this study, we use salivary glands and fat body cells of *Drosophila* larvae to study the effect of dPIP_4_K on insulin receptor activated, Class I PI_3_K signalling. We find that in *Drosophila* larval salivary gland cells, loss of *dPIP_4_K* enhanced the growth-promoting effects of overexpressing components of the insulin signalling pathway. dPIP_4_K regulates the levels of PIP_3_ and the intrinsic sensitivity to insulin at the plasma membrane. Insulin signalling activity is regulated through negative feedback from TORC_1_ in cells [27,28]. This TORC_1_ dependent reduction in insulin-stimulated PIP_3_ production is rendered ineffective in the absence of dPIP_4_K. Finally, we show that these cellular changes in insulin signalling have consequences on circulating sugar metabolism in larvae and also their susceptibility to insulin resistance on a high-sugar diet. Altogether, we demonstrate an important physiological role for dPIP_4_K as a negative regulator of Class I PI_3_K signaling during insulin stimulation in *Drosophila in vivo*.

## Results

### dPIP_4_K genetically interacts with the insulin receptor signalling pathway

Salivary glands are endo-replicative organs in *Drosophila* larvae that are composed of large polarized polyploid cells. Previously, we have demonstrated the use of this organ as a model to study changes in cell size [11,26]. Prior studies on insulin receptor signalling (scheme depicted in Fig. 1A) have revealed a role for this pathway in the autonomous control of both cell size and proliferation [23,25]. However, direct evidence for such regulation in salivary glands has not been demonstrated. Therefore, as a proof of principle, we depleted *dInR* levels through RNA interference (RNAi) selectively in the salivary glands of 3^rd^ instar larvae using the driver *AB1*Gal_4_. As expected, this resulted in a reduction of the average size of salivary gland cells without a change in the number of cells (Fig. 1B, C and S1A). Likewise, overexpression of *dInR* (Fig. 1D(i) and S1B) and *chico* (Fig. 1E(i) and S1D) selectively in the salivary glands also results in an increase in cell size. Thus, insulin receptor signalling regulates cell size in the salivary gland.

**Fig. 1.**
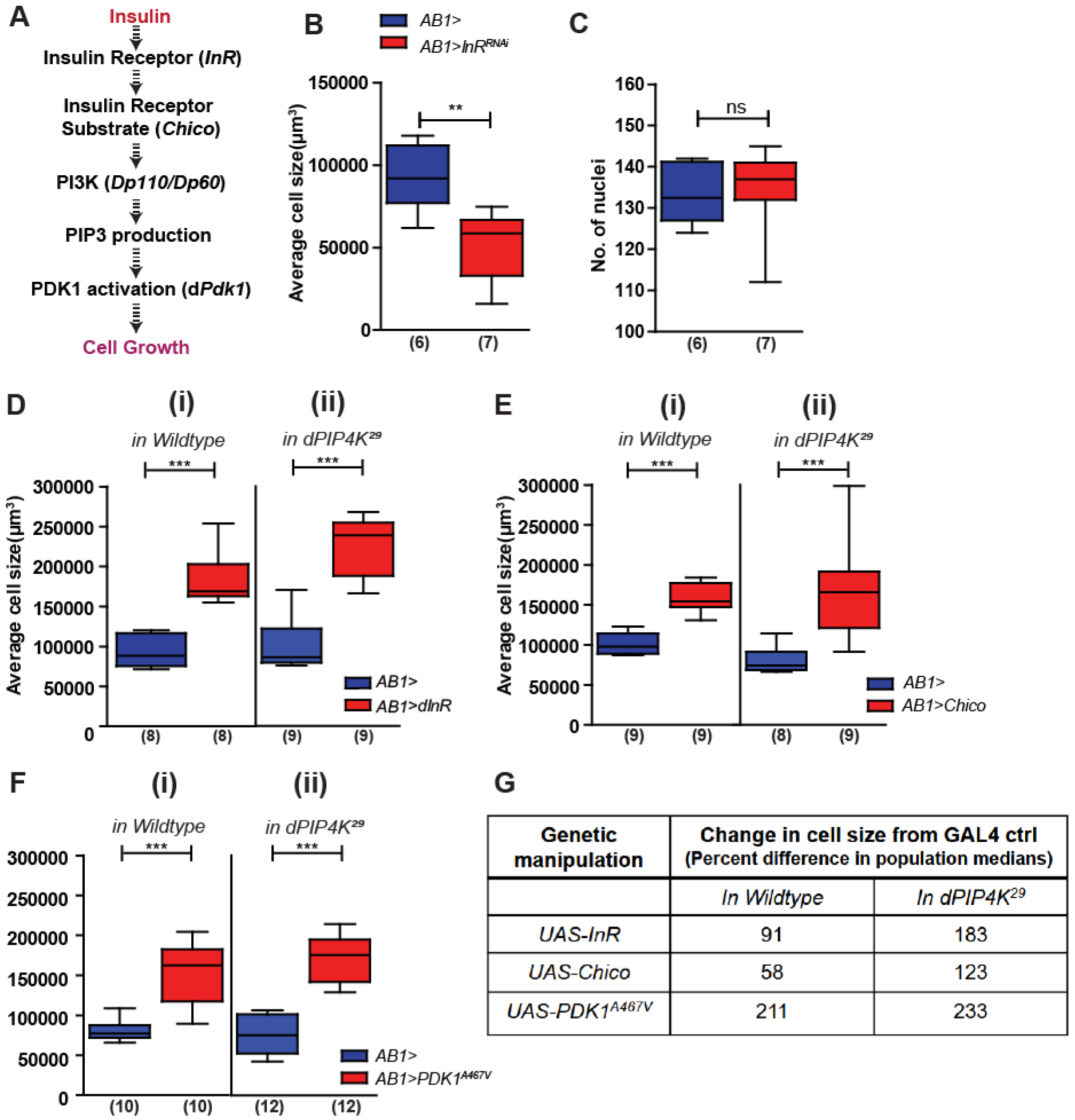
dPIP_4_K epistatically interacts with insulin receptor signalling to modulate cell size. **A.** Schematic depicting the components of insulin signalling cascade studied in the subsequent experiments. **B-G.** Salivary gland cell size measurements. **B.** Cell sizes upon knockdown of Insulin receptor in salivary glands. **C.** Quantification of the no. of nuclei. Whiskers in the box plots represent minimum and maximum values, with a line at the median. Cell size measurements in wildtype (*ROR*) and *dPIP_4_K*^*29*^ backgrounds as indicated upon - **D.** Overexpression of insulin receptor (InR) **E.** Overexpression of insulin receptor substrate - Chico **F.** Overexpression of PDK1^A467V^ **G.** Table with differences in median values of cell sizes across different genetic manipulations. Numbers inside the parentheses below the plots indicate the no. of biological replicates used for the measurement. *Mann Whitney test* used for statistical analysis of the distributions. \*\**p-value* <0.01, \*\*\**p-value* >0.001. Genotypes: **B, C.** *AB1Gal_4_/*+ *and AB1Gal_4_; UAS-dINR*^*RNAi*^. **D(i).** *AB1Gal_4_*/+ *and UAS-dINR/+; AB1Gal_4_/+* and **(ii).** *AB1Gal_4_*/+; *dPIP_4_K*^*29*^ *and UAS-dINR/+; AB1Gal_4_/+; dPIP_4_K*^*29*^. **E(i).** *AB1Gal_4_*/+ *and UAS-Chico*/+; *AB1Gal_4_*/+ and **(ii).** *AB1Gal_4_/+; dPIP_4_K*^*29*^ *and UAS-Chico*/+; *AB1Gal_4_*/+; *dPIP_4_K*^*29*^. **F(i).** *AB1Gal_4_*/+ and *UAS-PDK*^*A467V*^/+; *AB1Gal_4_*/+ and **(ii).** *AB1Gal_4_*/+; *dPIP_4_K*^*29*^ *and UAS-PDK*^*A467V*^/+; *AB1Gal_4_*/+; *dPIP_4_K*^*29*^.

We then compared the effect of overexpressing *dInR* in wild-type and *dPIP_4_K*^*29*^ cells. When *dInR* was over-expressed in *dPIP_4_K*^*29*^ (*AB1>dInR*; *dPIP_4_K*^*29*^), we also found an increase in salivary gland cell size (Fig.1D(ii) and S1C); but the increase in cell size elicited was significantly greater than that seen in wild-type cells (*AB1>dInR*) (Fig. 1G). Similar results were seen when comparing the effect of *chico* overexpression in wild-type and *dPIP_4_K*^*29*^ cells; i.e. *chico* overexpression elicited a larger increase in cell size in *dPIP_4_K*^*29*^ compared to wild-type (Fig. 1E(ii) and Fig. S1E).

Further, in order to decipher how dPIP_4_K specifically interacted with the insulin signalling pathway we tested if constitutively activating a downstream step will abolish the differences between wild-type and *dPIP_4_K*^*29*^ cells. For this, we expressed a constitutively active form of <u>P</u>hosphoinositide-<u>D</u>ependent *K*inase-1 (PDK1) (PDK1^A467V^) which is normally activated by PIP_3_ downstream of insulin receptor activation and regulates cell growth [29,30]. Expression of PDK1^A467V^ in salivary glands results in an increase in cell size (Fig. 1F(i) and S1F) and this was also seen when PDK1^A467V^ was expressed in *dPIP_4_K*^*29*^ (Fig. 1F(ii) and S1G). However, in contrast to *dInR* and *chico* manipulations, the effect of overexpressing PDK1^A467V^ resulted in an equivalent cell size increase in wild-type and *dPIP_4_K*^*29*^ (Fig. 1G). These findings suggest that in *Drosophila* larval cells dPIP_4_K modulates insulin receptor signalling at a step likely prior to PDK1 activation.

### PIP_3_ levels are elevated in dPIP_4_K depleted larval tissues

An essential early event in InR signal transduction is the activation of Class I PI_3_K leading to the production of PIP_3_ at the plasma membrane [1]. Therefore, we measured PIP_3_, levels at the plasma membrane by imaging salivary glands from wandering third instar larvae expressing a PIP_3_-specific probe (GFP::PH-GRP1) [31]. We observed that under basal conditions, plasma membrane PIP_3_ in *dPIP_4_K*^*29*^ showed a small but significant elevation compared to wild-type cells (Fig. 2A(i) and (ii)). Similar results were observed in experiments with fat body cells, i.e. PIP_3_ levels in *dPIP_4_K*^*29*^ fat body cells were elevated compared to wild type (Fig. 2B(i) and (ii)).

**Fig. 2.**
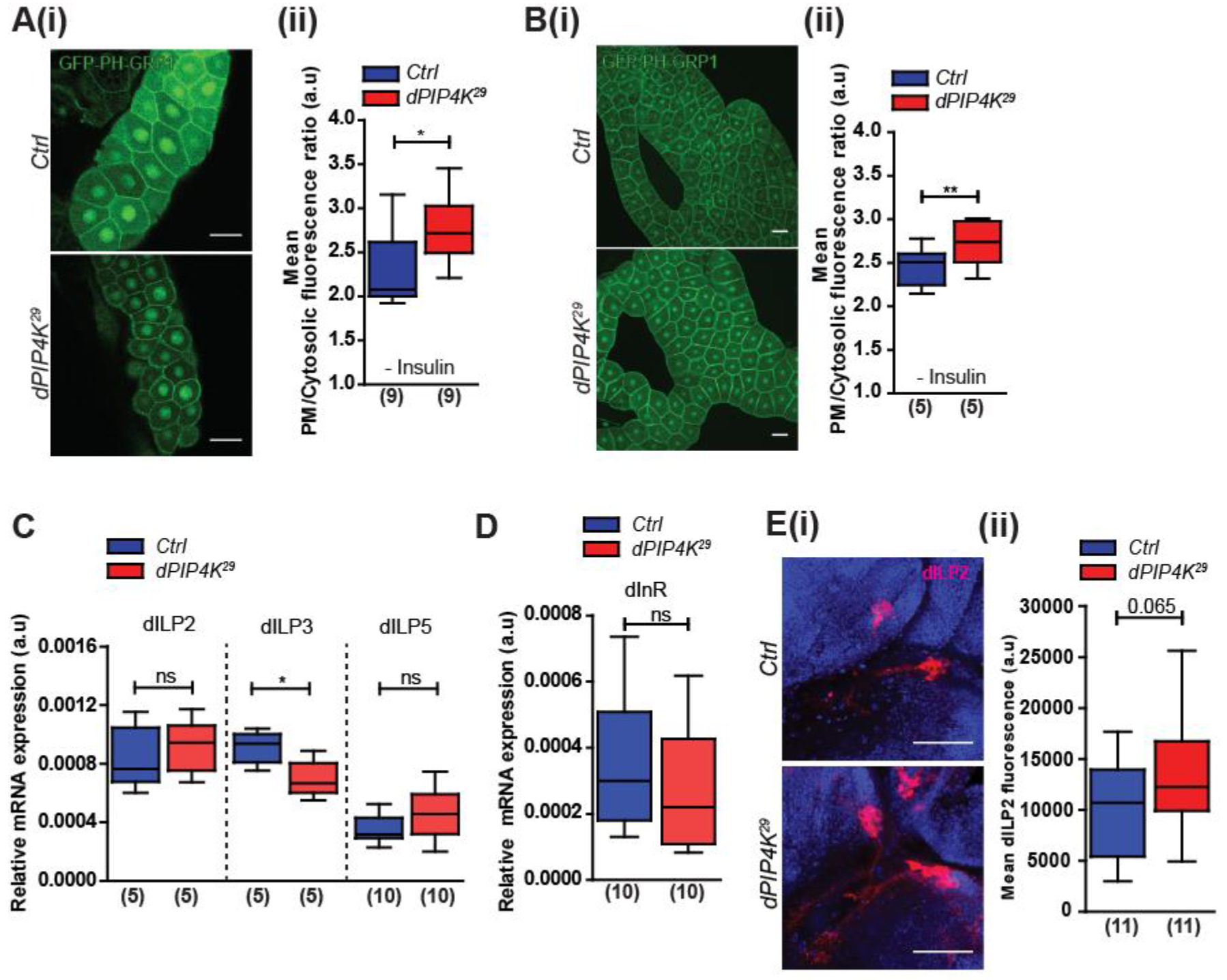
Plasma membrane PIP_3_ levels are elevated in *dPIP_4_K* mutant tissues without an increase in humoral dILP secretion. **A-B.** PIP_3_ quantification - Images showing the intensity and localization of the PIP_3_ binding probe (GFP-PH-GRP1) in larval tissues. The PIP_3_ levels quantified as the ratio of the probe fluorescence intensity on the plasma membrane to that in the cytosol. **A(i).** Representative confocal images showing the distribution of the probe in the salivary glands. **A(ii).** Quantification of PIP_3_ levels between control and *dPIP_4_K*^*29*^ salivary glands for experiment depicted in A(i). (Mean fluorescence intensity ratios calculated from a minimum of 10 cells from each salivary gland). **B(i).** Representative confocal images showing the distribution of the probe in the fat body and in **B(ii)** the quantification from these experiments. The ratio was calculated from about 50 cells from 12 fat body regions pooled from 5 animals for each genotype respectively. Wandering third instar larvae used for measurements. Scale: 50 μm for salivary gland images and 10 μm for fat body images C. qPCR measurements for mRNA levels of *dILP 2, 3* and *5* from whole larvae. Transcript levels for each gene were normalized to the mRNA levels of *rp49* in the same sample. **D.** qPCR measurements for *dInR*. Transcript levels for each gene were normalized to the mRNA levels of *rp49* in the same sample **E(i).** Confocal z-projections showing immunostaining for dILP2 in larval IPCs, Scale: 50 μm. **E(ii).** Quantification of dILP2 staining intensity in the third instar wandering larval brains. Whiskers in the box plots represent minimum and maximum values, with a line at the median. Numbers inside the parentheses below the plots indicate the no. of biological replicates used for the measurement. *Mann Whitney test* used for statistical analysis of the distributions. \**p-value* < 0.05, \*\**p-value* <0.01. Genotypes: **A-D.** *tGPH and tGPH; dPIP_4_K*^*29*^. **E.** *ROR and dPIP_4_K*^*29*^.

During larval development in *Drosophila*, nutritional cues and other signals result in the release of *<u>D</u>rosophila <u>I</u>nsulin*-*<u>l</u>ike <u>p</u>eptides* (dlLPs) [32] that bind to and activate dInR triggering Class I PI_3_K activation and PIP_3_ production. The elevated PIP_3_ levels observed in *dPIP_4_K*^*29*^ tissues could, therefore, result from (i) enhanced production and release of dlLPs (ii) upregulation in insulin receptor levels (iii) increase in activity of insulin receptor or events downstream of receptor activation. To distinguish between these possibilities, we performed Q-PCR analysis to measure the levels of *dILP2, 3, 5* mRNAs [the levels of these are known to be transcriptionally regulated] [23]. We found that the transcript levels for these dlLPs were not upregulated in *dPIP_4_K*^*29*^ compared to wildtype (Fig. 2C). Since these were not altered, we tested for enhanced dILP release, by measuring the levels of dILP2 within the neurosecretory insulin-producing cells (IPCs) from the brains of wandering third instar larvae. Immunoreactivity for dILP2 produced in IPCs is expected to be lower when more of it is released into the hemolymph. We found that the average intensity of dILP2 immunostaining in the IPCs was not lower in *dPIP_4_K^29^* compared to controls; instead, it showed a small increase (Fig. 2E(i) and (ii)). Thus, we found no evidence of elevated production and release of dlLPs in 3^rd^ instar larvae that might explain the increased PIP_3_ levels observed in *dPIP_4_K*^*29*^. Further, we observed that *InR* receptor mRNA levels were also not different between *dPIP_4_K*^*29*^ and wildtype indicating that levels of InR that are activated by dlLPs are also not likely to be different between the two genotypes (Fig. 2D). Collectively, our experiments show plasma membrane PIP_3_ levels to be elevated in cells lacking dPIP_4_K without an increase in dILP secretion or cellular insulin receptor levels.

### *dPIP_4_K*^*29*^ cells are intrinsically more sensitive to insulin stimulation

We developed *ex-vivo* assays to test the sensitivity of larval tissues to bovine insulin. It has previously been shown that *Drosophila* cells respond to bovine insulin using signal transduction elements conserved with those proposed for the canonical mammalian insulin signalling pathway [33]. We observed that in salivary glands and fat body dissected from 3^rd^ instar larvae, *ex-vivo* insulin stimulation triggered a rise in plasma membrane PIP_3_ levels, measured using the GFP::PH-GRP1 probe. Interestingly, following insulin stimulation (10 min, 10 μM), the rise in PIP_3_ levels in *dPIP_4_K*^*29*^ was higher than in wild type (Fig. 3A(i), A(ii)). Selective depletion of dPIP_4_K protein using RNAi specifically in the salivary gland cells also produced a similar result. (Fig. 3C(i) and (ii)). The increased sensitivity of *dPIP_4_K^29^* cells to *ex-vivo* insulin stimulation could be reverted by specifically reconstituting dPIP_4_K in salivary gland cells (Fig. 3D). Overexpression of dPIP_4_K in wild-type salivary gland cells resulted in reduced levels of insulin-stimulated PIP_3_ levels (Fig. 3E). A similar observation was made in fat body cells dissected from larvae where PIP_3_ production increased with stimulation over a wide range (100-fold) of insulin concentrations used. While 100 nM of insulin barely elicited an increase in plasma membrane PIP_3_ levels, we observed that *dPIP_4_K*^*29*^ fat cells show a larger rise in PIP_3_ levels compared to the controls at higher concentrations (Fig. 3C(i)-(iii)).

**Fig. 3.**
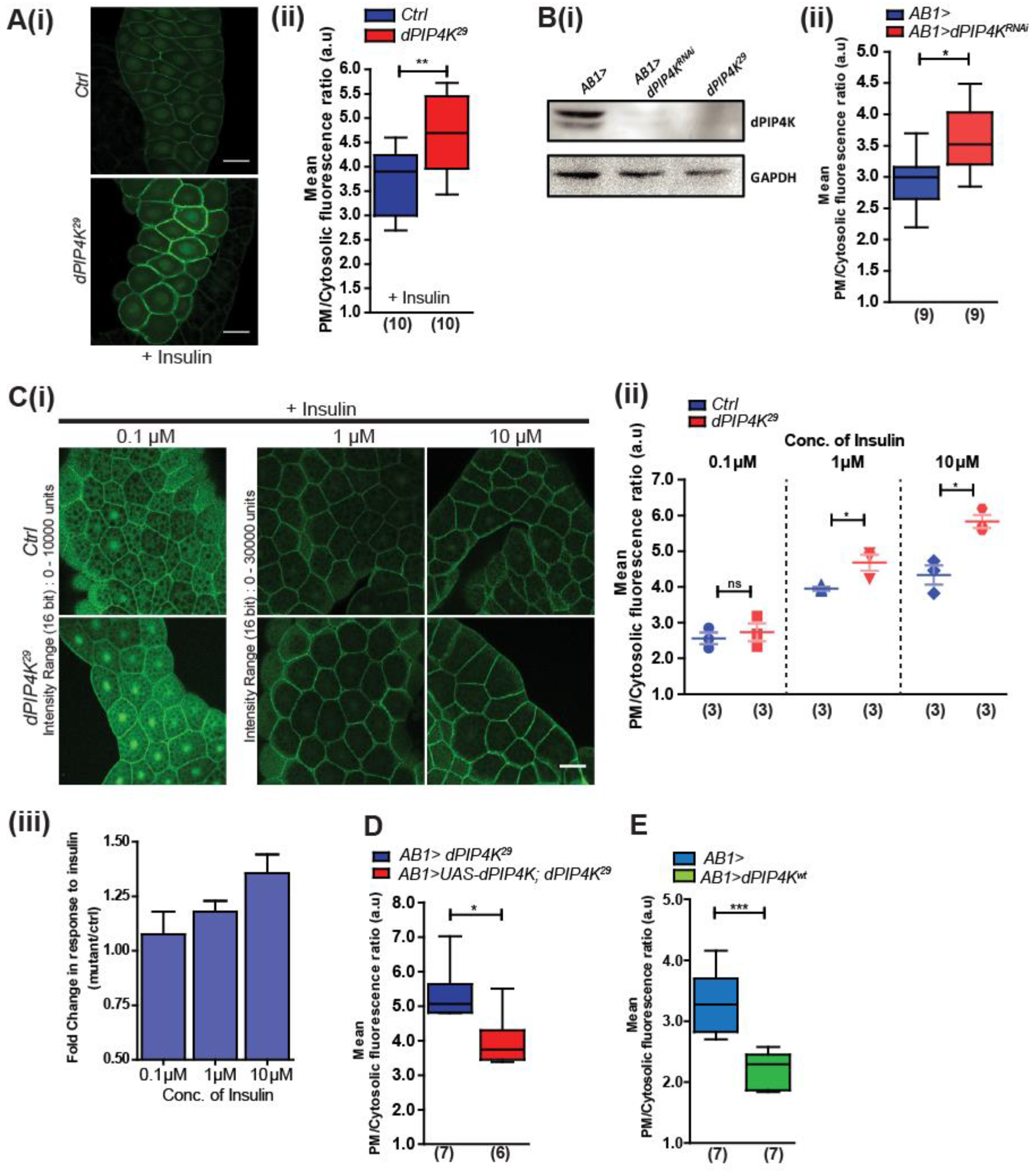
Increased sensitivity of *Drosophila* larval cells to insulin upon loss of dPIP_4_K. **A(i).** Confocal z-projections showing levels and localization of the PIP_3_ probe in control and *dPIP_4_K*^*29*^ in salivary glands cells stimulated with 10 μM insulin for 10 min and in **A(ii)**, quantification of PIP_3_ levels between control and *dPIP_4_K*^*29*^ salivary glands from the same set of experiments. Immunoblot (from wandering third instar stage) showing a reduction in levels of dPIP_4_K protein in salivary gland lysates upon knockdown of dPIP_4_K using *AB1*Gal_4_. **B(ii).** Relative quantification of PIP_3_ levels using the GFP-PH-GRP1 probe between control and *dPIP_4_K*-knockdown salivary glands **C(i).** Confocal z-projections of fat body lobes expressing PIP_3_ binding probe from control and *dPIP_4_K*^*29*^ late third instar larvae stimulated with 0.1 μM, 1 μM and 10 μM insulin post 2 hr starvation. **C(ii).** Quantification of PIP_3_ for experiments in fig. C (50 cells from at least 3 samples in the fat body for each genotype and treatment used for analysis). Scale: 50 μm. **C(iii).** Comparison of mean fold change (mutant w.r.t control) in response to insulin computed from data in **B(ii).** Comparison of PIP_3_ levels between **(D)** mutant and rescue salivary glands and **(E)** wildtype and salivary glands overexpressing dPIP_4_K. Whiskers in the box plots represent minimum and maximum values, with a line at the median. Numbers inside the parentheses below the plots indicate the no. of biological replicates used for the measurement. Scale: 50 μM. *Mann Whitney test* used for statistical analysis of the distributions. \**p-value* <0.05, \*\**p-value* <0.01, \*\*\**p-value* <0.001 Genotypes: **A, C.** *tGPH and tGPH; *dPIP_4_K*^*29*^. **B.** *AB1Gal_4_, tGPH*/+ and *dPIP_4_K*^*RNAi*^/+; AB1Gal_4_, tGPHI+.* **D.** *AB1Gal_4_, tGPH*/+; *dPIP_4_K*^*29*^ and *UAS-dPIP_4_K/+; AB1Gal_4_, tGPH/+; dPIP_4_K*^*29*^. **E.** *AB1Gal_4_, tGPH/+* and *UAS-dPIP_4_K/+; AB1Gal_4_, tGPH/+.*

### Quantitative measurements of PIP_3_ mass in *Drosophila* larvae

To test if the probe-based imaging of PIP_3_ in single cells indeed reflects *in vivo* changes across the animal, we refined and adapted existing protocols [34] to perform mass spectrometric measurements of PIP_3_ from *Drosophila* whole larval lipid extracts (Scheme depicted in Fig. 4A). The amount of PIP_3_ that has been detected and quantified from biological samples is in the range of a few tens of picomoles [35]. We coupled liquid chromatography to high sensitivity mass spectrometry (LCMS) and used a Multiple Reaction Monitoring (MRM) method to detect PIP_3_ standards for reliable quantification down to a few femtomoles (ca. 10 fmol, the lowest point in the figure inset on the standard curve in Fig. S2(A). Since cellular lipids are composed of molecular species with varying acyl chain lengths, we first characterized the PIP_3_ species from *Drosophila* whole larval extracts through use of neutral loss scans and thereafter quantified the abundance of these species. Fig. S2(B) depicts the elution profiles of the different PIP_3_ species that were reproducibly detected across samples and Fig. S3 (A) shows the relative abundance of various PIP_3_ species. The 34:2 PIP_3_ species was found to be the most abundant. To standardize the procedure, we bisected whole larvae, stimulated them with insulin and measured the levels of various PIP_3_ species between samples with and without insulin stimulation. Our LCMS method could clearly detect an increase in the levels of several PIP_3_ species upon insulin stimulation (Fig. S3(B)).

**Fig. 4.**
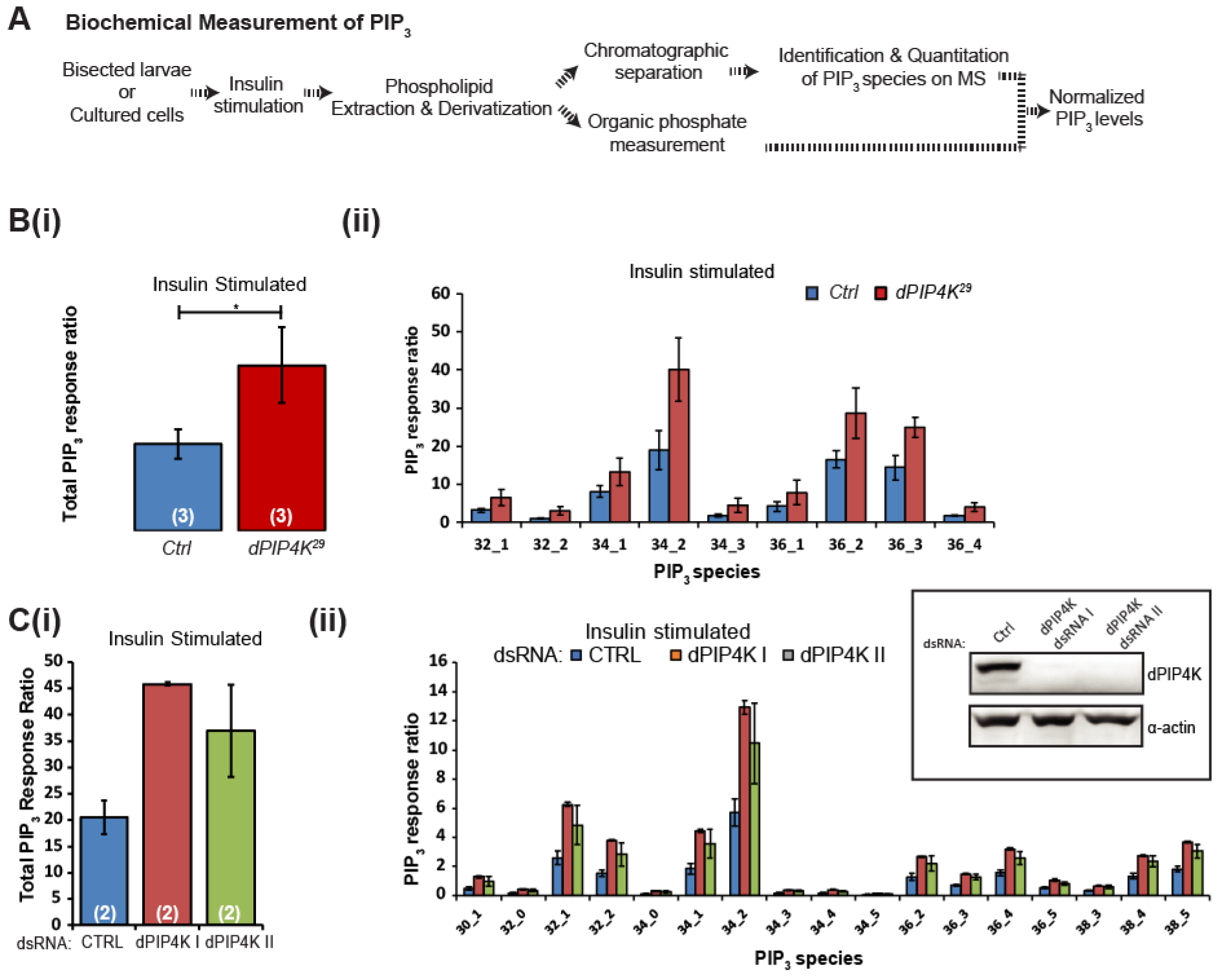
Quantitative biochemical measurements of PIP_3_ identify dPIP_4_K as a negative regulator of insulin signalling. **A.** Schematic outline of the steps involved in LCMS-based measurement of PIP_3_ lipid from larvae and cells upon insulin stimulation. **B(i).** Measurement of total PIP_3_ levels in whole larval lipid extracts from wildtype (*ROR*) and *dPIP_4_K*^*29*^ using LCMS. **B(ii).** Levels of various larval PIP_3_ species in wildtype (*ROR*) and *dPIP_4_K*^*29*^ whole larval lipid extracts. **C(i).** Measurement of total PIP_3_ levels using LCMS in whole cell lipid extracts from S2R+ cells treated with indicated dsRNAs. **C(ii).** Levels of various larval PIP_3_ species in whole cell lipid extracts from S2R+ cells. The graphs show mean PIP_3_ levels (normalized to spiked internal standards and total organic lipid phosphates recovered). Error bars depict SD. Inset **C(ii).** Immunoblot showing the knockdown of dPIP_4_K in S2R+ cells using two different sets of dsRNAs. *Student’s unpaired t-test* used for statistical analysis of the distributions, \**p-value* < 0.05. On each graph, numbers inside the parentheses indicate the no. of biological replicates used for the measurement.

Using this method, we compared PIP_3_ levels from whole larval lipid extracts of various genotypes following insulin stimulation. We observed that compared to controls, (*dPIP_4_K*^*29*^ larvae showed higher PIP_3_ levels upon insulin stimulation (Fig. 4B(i), (ii)). Similarly, upon pan-larval knockdown of dPIP_4_K by RNAi enhanced PIP_3_ levels were observed following insulin stimulation (Fig. S3C(ii) and (iii)) although the differences were not as striking as seen in *dPIP_4_K*^*29*^; presumably this reflects the residual and variable amounts of dPIP_4_K protein seen during RNAi based knockdown ((Fig. S3C(i)). We also performed pan-larval rescue of dPIP_4_K protein in *dPIP_4_K*^*29*^ larvae and observed a trend of rescue in levels of various PIP_3_ species (Fig. S3D(i) and (ii)). Finally, in an alternate setting, we also depleted dPIP_4_K in *Drosophila* S2 cells (inset in Fig. 4C(ii)) in culture using two independent dsRNA treatments and found that on insulin stimulation of serum starved cells, the total level of PIP_3_ was enhanced compared to that in control cells (Fig. 4C(i)); the levels of individuals species of PIP_3_ underlying this elevation broadly reflected those seen in experiments with *Drosophila* larval extracts (Fig. 4C(ii)). Together, the observations from these two independent assays (fluorescent probe based PIP_3_ measurement and mass spectrometry) suggests that in dPIP_4_K depleted cells, increased amounts of PIP_3_ are produced at the plasma membrane during insulin stimulation, thus implying that *dPIP_4_K* negatively regulates PIP_3_ production.

### dPIP_4_K directly regulates insulin receptor signalling independent of TORC_1_

We had previously reported a systemic reduction in TORC_1_ activity in *dPIP_4_K*^*29*^ larvae [11]. It is reported in mammalian cells that TORC_1_ activity can mediate feedback inhibition on insulin receptor substrate (IRS) to suppress insulin signalling and conversely reduced TORC_1_ activation can increase insulin signaling [28,36,37](Proposed feedback depicted in Fig.5 scheme 1 and 2). We tested if the increased insulin-stimulated plasma membrane PIP_3_ levels were a result of reduced cellular TORC_1_ activation in *dPIP_4_K*^*29*^. We down-regulated Rheb, the GTPase that directly binds and activates TORC_1_ [38] in the salivary gland. *AB1 >Rheb^BNAi^* glands have substantially reduced cell size consistent with the known requirement for TORC_1_ signalling in regulating cell size (Fig. 5 A (i)). Following insulin stimulation of *AB1>dRheb*^*RNAi*^ glands, PIP_3_ levels at the plasma membrane were elevated compared to controls (Fig. 5A (ii)). Conversely, we compared PIP_3_ production in control cells and those selectively overexpressing Rheb (*AB1>dRheb*) that is expected to enhance TORC_1_ activity. Following insulin stimulation, the levels of PIP_3_ generated were significantly lower in *AB1 >dRheb* glands compared to controls (Fig. 5B (i), (ii)). Similarly, knockdown of TSC, the GTPase activating protein (GAP) for Rheb, expected to result in hyperactivation of Rheb [39], also reduces the PIP_3_ levels seen post insulin stimulation (Fig. 5C (i), (ii)). These results demonstrate that TORC_1_ output can control plasma membrane PIP_3_ levels during insulin signaling in salivary gland cells (Fig.4 scheme 1 and 2).

**Fig. 5.**
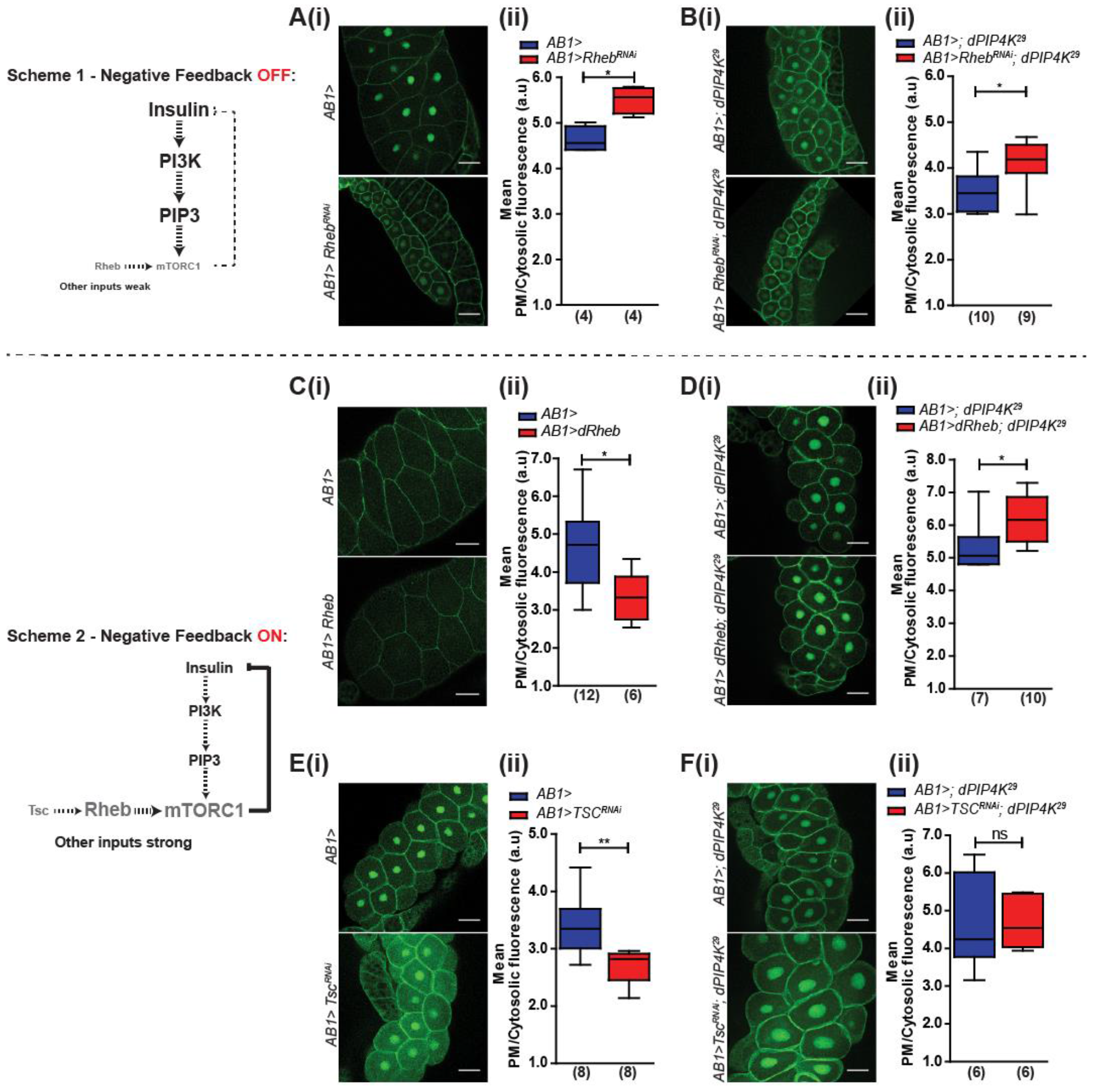
dPIP_4_K additionally regulates insulin signalling independent of TORC_1_-mediated negative feedback. **Scheme 1** and **2** depict the feedback regulation of insulin signalling by TORC_1_ activity in **OFF** and **ON** state respectively. The font and arrow sizes are indicative of the extent of molecular activity. **A-F.** PIP_3_ quantification ‒ Salivary gland images showing the distribution of the PIP_3_ binding probe GFP-PH-GRP1 in cells. The distribution was quantified as the ratio of probe fluorescence on the plasma membrane to that in the cytosol. In all these experiments, the genetic manipulation was restricted to the salivary glands using *AB1*Gal_4_. **A, C, E.** In a wildtype background, Downregulation of TOR signalling by RNAi for *Rheb* **[A (i and ii)].** Upregulation of TOR signalling through overexpression of *Rheb* **[C (i and ii)]**, knockdown of *Tsc* **[E (i and ii)] B, D, F.** In a *dPIP_4_K*^*29*^ background, Downregulation of TOR signalling by RNAi for *Rheb* **[B (i and ii)]** Upregulation of TOR signalling through overexpression ofRheb **[D (i and ii)]**, knockdown of Tsc **[F (iand ii)]**. Scale: 50 μM. Whiskers in the box plots represent minimum and maximum values, with a line at the median. Numbers inside the parentheses below the plots indicate the no. of biological replicates used for the measurement. *Mann Whitney test* used for statistical analysis of the distributions. \**p-value* < 0.05. \*\**p-value* <0.01. Genotypes: **A.** *AB1Gal_4_, tGPH*/+ *and AB1Gal_4_, tGPH*/ *UAS-Rheb*^*RNAi*^. **B.** *AB1Gal_4_, tGPH* /+; *dPIP_4_K*^*29*^ *and AB1Gal_4_, tGPH*/ *UAS-Rheb*^*RNAi*^; *dPIP_4_K*^*29*^. **C.** *AB1Gal_4_, tGPH*/+ *and UAS-dRheb*/+; *AB1Gal_4_, tGPH*/+. **D.** *AB1Gal_4_, tGPH*/+; *dPIP_4_K*^*29*^ *and UAS-dRheb*/+; *AB1Gal_4_, tGPH*/+; *dPIP_4_K*^*29*^. **E.** *AB1Gal_4_, tGPH*/+ *and UAS-Tsc1^mAi^/+; AB1Gal_4_, tGPH/+.* **F.** *AB1Gal_4_, tGPH /+; dPIP_4_K^29^ and UAS-Tsc1^RNAi^/+; AB1Gal_4_, tGPH/+; dPIP_4_K*^*29*^.

We tested the effect of dPIP_4_K function on TORC_1_-mediated control of PIP_3_ levels during insulin stimulation. When dPIP_4_K function is reconstituted in salivary glands (*AB1 >dPIP_4_K; dPIP_4_K*^*29*^), as expected, normal levels of insulin-stimulated PIP_3_ production were restored (refer Fig. 3D). Knockdown of *dRheb* in *dPIP_4_K*^*29*^ salivary glands reduced the size of cells as expected (Fig. S5B(ii)) but also resulted in a further elevation of insulin-stimulated PIP_3_ levels over that seen in *dPIP_4_K*^*29*^ (Fig. 5B(i), (ii)). However, when *dRheb* was overexpressed in *dPIP_4_K*^*29*^ salivary glands; (*AB1 >dRheb; dPIP_4_K*^*29*^), surprisingly, we found that insulin-stimulated PIP_3_ levels were not lower than in *AB1>; dPIP_4_K*^*29*^ (Fig. 5D (i), (ii)). Likewise, depletion of TSC in *dPIP_4_K*^*29*^ (*AB1>Tsc*^*RNAi*^; *dPIPK*^*29*^) did not lower insulin stimulated PIP_3_ levels (Fig. 5F (i), (ii)). Thus, enhanced activation of TORC_1_ in salivary glands failed to complement the elevated PIP_3_ levels resulting from loss of dPIP_4_K. Together, our findings indicate that dPIP_4_K function offers an additional mode of PIP_3_ regulation that acts independent of TOR1 activity during insulin signalling.

### PIP_4_K is required at the plasma membrane to control of insulin-stimulated PIP_3_ production

We and others have previously shown that PIP_4_KS localize to multiple subcellular membrane compartments [11,40]. It is also reported that the substrate for this enzyme i.e. PI_5_P is present on various organellar membranes inside cells [41]. To further probe the mechanism of regulation of PIP_3_ levels by dPIP_4_K, we decided to identify the sub-cellular compartment at which dPIP_4_K function is required to regulate PIP_3_ levels. We generated transgenic flies to target dPIP_4_K to specific subcellular compartments (Fig. 6A). Using unique signal sequences, we targeted dPIP_4_K specifically to the plasma membrane (Fig. 6B (ii)), endomembrane compartments viz. the ER and Golgi (Fig. 6B (iii)) and the lysosomes (Fig. 6B (iv)). Lysates from S2R+ cells expressing these constructs for assayed for PIP_4_K activity and we found that all of the targeted dPIP_4_K enzymes were active (Fig. 6C(i)-(ii)); activity was proportional to the amount of protein expressed. Each of these targeted dPIP_4_K constructs were selectively reconstituted into dPIP_4_K null (*dPIP_4_K*^*29*^) cells and tested for its ability to revert the enhanced insulin-stimulated PIP_3_ production of *dPIP_4_K*^*29*^. For this, we stimulated dissected salivary glands *ex-vivo* with insulin and measured PIP_3_, production using the GFP::PH-GRP1 probe. Under these conditions, while endomembrane (Fig. 6E) and lysosome-targeted (Fig. 6F) dPIP_4_K failed to revert the elevated PIP_3_ levels of *dPIP_4_K*^*29*^, reconstitution with the plasma-membrane targeted dPIP_4_K completely restored the elevated PIP_3_ levels in *dPIP_4_K*^*29*^ to that of controls (Fig. 6D). Further, overexpression of plasma-membrane targeted dPIP_4_K in wildtype salivary gland cells resulted in lower insulin stimulated PIP_3_ levels compared to the controls at 5 mins post insulin stimulation (Fig. 6G). These observations suggest that plasma membrane localized dPIP_4_K is sufficient to regulate insulin-stimulated PIP_3_ production.

**Fig. 6.**
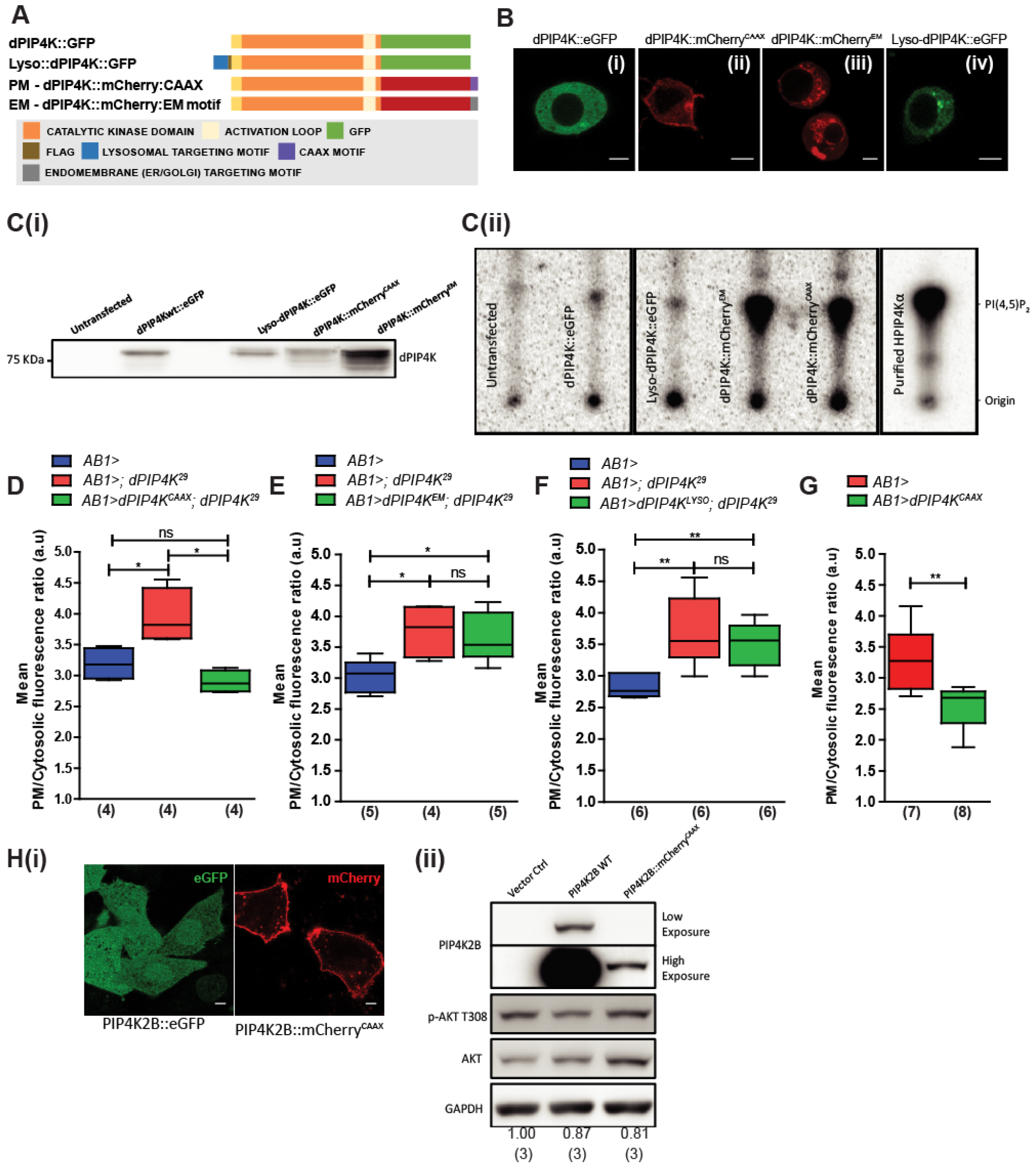
PIP_4_K functions at the plasma membrane as a negative regulator of insulin receptor signalling. **A.** Schematic showing constructs that target dPIP_4_K to different subcellular compartments and the motifs used for targeting. **B (i-iv).** Representative confocal z-projections of S2R+ cells with act-Gal_4_ expressing various dPIP_4_K constructs **(i)** wildtype GFP-tagged dPIP_4_K **(ii)** plasma-membrane (PM) targeted mCherry-tagged dPIP_4_K **(iii)** mCherry tagged dPIP_4_K targeted to various intracellular membranes inclusive of ER, Golgi and endo-lysosomal system **(iv)** GFP-tagged dPIP_4_K targeted to the lysosome. **C(i).** Immunoblots from S2R+ lysates showing the expression of indicated dPIP_4_K constructs that were used in the *in vitro* assay. **C(ii).** *In vitro* kinase assay from S2R+ cell lysates showing the activity of different overexpressed dPIP_4_K constructs. PIP_3_ measurement on insulin stimulation (10 μM) using the PH-GFP-GRP1 probe **D-F.** in *dPIP_4_K*^*29*^ salivary glands reconstituted with **D.** PM targeted dPIP_4_K, **E.** endo-membrane targeted dPIP_4_K, **F.** Lysosomal dPIP_4_K, **G.** upon overexpression of dPIP_4_K::mCherry^CAAX^ in the salivary glands. Whiskers in the box plots represent minimum and maximum values, with a line at the median. Numbers inside the parentheses below the plots indicate the no. of biological replicates used for the measurement. *Mann Whitney test* used for statistical analysis of the distributions. \**p-value* < 0.05. \*\**p-value* <0.01. **H(i).** Representative confocal z-projections of CHO-IR cells overexpressing GFP-PIP_4_K_2_B and PIP_4_K_2_B::mCherry-CAAX. **H(ii).** Immunoblots from CHO-IR cells expressing PIP_4_K_2_B constructs stimulated with 1 μM insulin for 10 min. The values below the blots represent the mean pAKT/Total AKT ratio across three independent experiments. Genotypes: **D.** *AB1Gal_4_, tGPH*/+ *and AB1Gal_4_, tGPH*/+; *dPIP_4_K*^*29*^ *and AB1Gal_4_, tGPH* / dPIP_4_K::mCherryCAAX; dPIP_4_K^29^. **E.** *AB1Gal_4_, tGPH I + and AB1Gal_4_, tGPH /+; dPIP_4_K^29^ and AB1Gal_4_, tGPH /dPIP_4_K::mCherryEM; dPIP_4_K^29^. **F.** *AB1Gal_4_, tGPH* /+ *and AB1Gal_4_, tGPH*/+; *dPIP_4_K*^*29*^ *and AB1Gal_4_, tGPH/Lysosomal-dPIP_4_K::eGFP; dPIP_4_K**^*29*^.

We also tested the ability of plasma membrane localized PIP_4_K to regulate steps downstream of PIP_3_ production during insulin signalling. In a previous study, overexpression of human PIP_4_K_2_B in CHO-IR cells was shown to reduce the levels of pAKT^T308^, an important PIP_3_ dependent signalling event during insulin stimulation [16]. We tested the effect of over expressing plasma membrane restricted PIP_4_K_2_B in these cells on pAKT^T308^ during insulin stimulation. We generated a PIP_4_K_2_B construct with a CAAX-motif at its C-terminus (PIP_4_K_2_B::mCherry^CAAX^) that localized the enzyme to the plasma membrane as expected, while the wildtype PIP_4_K_2_B (PIP_4_K_2_B::eGFP) can be seen at various subcellular compartments (Fig. 6H(i)). CHO-IR cells transiently overexpressing either PIP_4_K_2_B::eGFP or PIP_4_K_2_B::mCherry^CAAX^ were serum starved, stimulated with insulin and pAKT^T308^ was measured through immunoblotting. As previously reported, we found that PIP_4_K_2_B::eGFP overexpression resulted in a small but significant decrease in pAKT^T308^ (Fig. 6H(ii)). Interestingly, consistent with our findings in *Drosophila* larval cells, we observed that over-expressed PIP_4_K_2_B::mCherry^CAAX^ also caused a decrease in pAKT^T308^. In fact, this decrease was achieved despite lower levels of expression of PIP_4_K_2_B::mCherry^CAAX^ compared to the wildtype protein. Thus, PIP_4_K_2_B activity at the plasma membrane is sufficient to negatively regulate PIP_3_ dependent pAKT^T308^ levels in mammalian cells.

### dPIP_4_K alters PIP_3_ turnover by modifying Class I PI_3_K activity

PIP_3_ levels at the plasma membrane upon insulin stimulation also depend on the length of time the receptor remains activated. Our 10-min stimulation protocol was based on earlier studies performed on *Drosophila* S2 cell-cultures where the response to insulin was maximal at 10 min [33]. However, in order to check for any differences in the dynamics of response to insulin, we also studied the time course of PIP_3_ elevation following increasing times of insulin stimulation *ex-vivo*. Comparison of fixed preparations of salivary glands expressing GFP-PH-GRP1 probe showed a comparable time course of PIP_3_ elevation but higher PIP_3_ levels at every time point in *dPIP_4_K*^*29*^ than in control glands (Fig. S4(i) and (ii)). To understand the effect of dPIP_4_K on insulin signaling at the plasma membrane with increased temporal resolution, we developed a live-imaging assay to follow the dynamics of PIP_3_ turnover using the PH-GRP1 probe in salivary gland cells. A schematic of the reactions involved in the process and the assay protocol is depicted in Fig. 7A(i) and (ii). In this assay, during insulin stimulation, the dynamics of PIP_3_ turnover has three phases ‒ (i) Rise phase ‒ PIP_3_ levels increase after a stimulus owing to the activation of PI_3_K and relatively lower phosphatase activity (ii) Steady-state phase ‒ the opposing kinase and phosphatase activities regulating PIP_3_ levels balance out each other (iii) Decay phase ‒ PI_3_K activity is irreversibly inhibited by wortmannin while PIP_3_ phosphatases remain active. A single experimental trace is shown in Fig. 7B (i); insulin stimulation triggers a rise in PIP_3_ levels that peak and subsequently decline. Addition of wortmannin prior to addition of insulin abolished insulin stimulated PIP_3_ production establishing the effectiveness of Class I PI_3_K inhibition in this assay (Fig. 7B(ii)). Under similar conditions, addition of the DMSO vehicle post insulin did not reduce PIP_3_ levels (Fig. 7B(iii)).

**Fig. 7.**
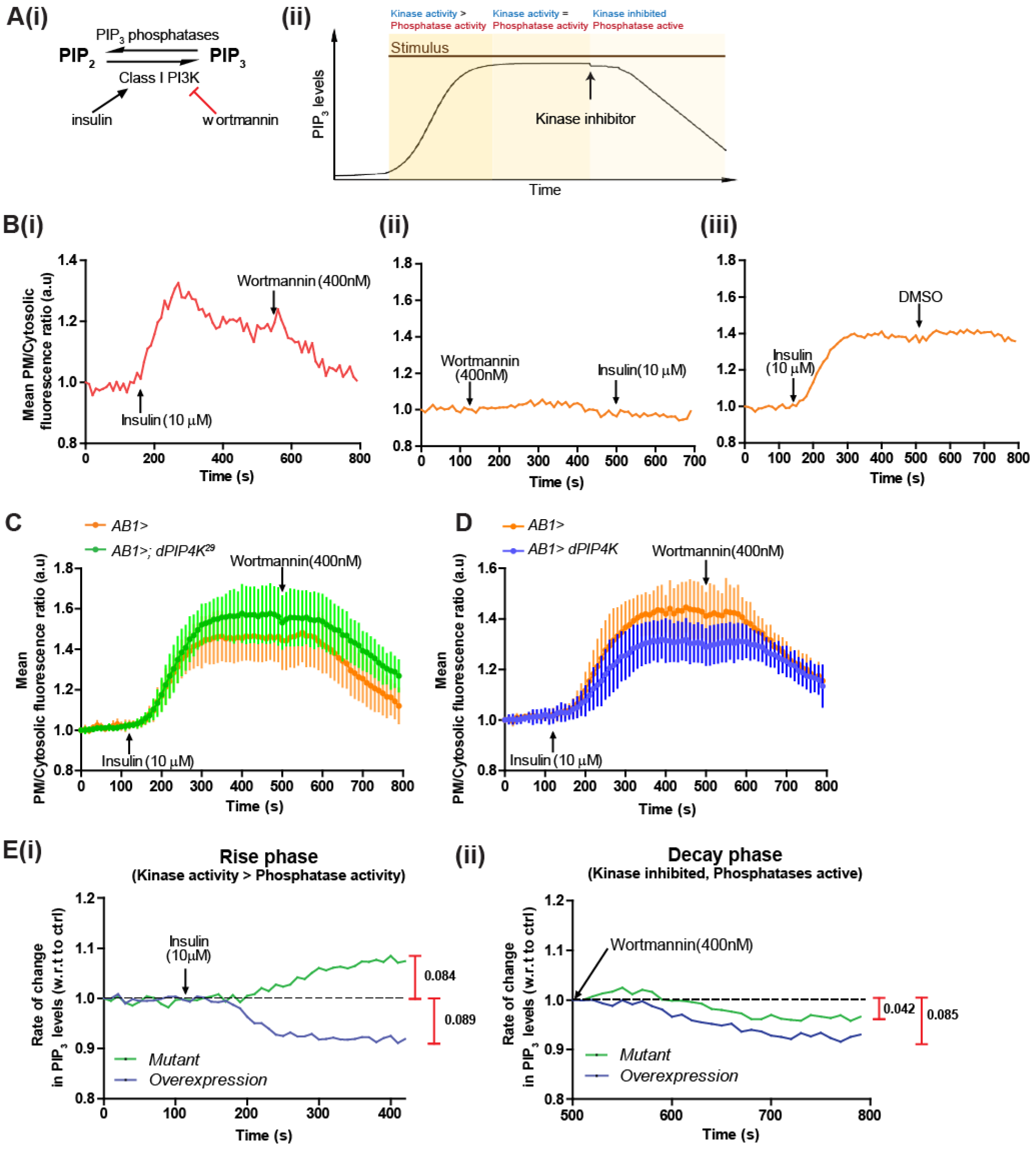
dPIP_4_K influences PIP_3_ turnover. **(A(i))** Schematic of the reactions that determine PIP_3_ turnover at the plasma membrane. Insulin stimulates PI_3_K activation. Wortmannin irreversibly inhibits PI_3_K activity. **(A(ii))** Live imaging assay protocol to follow PIP_3_ dynamics with three phases as depicted. **(B(i))** A single trace from live imaging of salivary glands expressing GFP-PH-GRP1 probe showing the changes in the plasma-membrane to cytosolic ratio of the probe fluorescence over time. **(B(ii))** Wortmannin addition (400nM) completely blocks insulin (10μM) induced PIP_3_ production **(B(iii))** DMSO (vehicle) addition does not reduce PIP_3_ levels post-insulin (10μM) stimulation. **(C)** and **(D)** Comparison of average traces of GFP-PH-GRP1 fluorescence ratios from multiple imaging runs of *control, dPIP_4_K*^*29*^ and *dPIP_4_K overexpression* salivary glands (N=7 for all genotypes). Error bars indicate SD. The two experiments were performed at different times, hence controls samples were repeated. **(E(i) and (ii))** Curves depict changes in slopes of fluorescence calculated by taking ratios of fluorescence from test genotypes to that in controls. The maximal difference is indicated alongside the graph. Genotypes: **B(i-iii).** *AB1Gal_4_, tGPH*/+. **C.** *AB1Gal_4_, tGPH*/+ and *AB1Gal_4_, tGPHI*+; *dPIP_4_K*^*29*^. **D.** *AB1Gal_4_, tGPH*/+ and *UAS-dPIP_4_K*/+; *AB1Gal_4_, tGPH*/+.

We tested the effect of loss of dPIP_4_K and tissue-specific overexpression of dPIP_4_K on PIP_3_ turnover. Loss of dPIP_4_K resulted in higher steady state levels of PIP_3_ in salivary gland cells compared to controls (Fig. 7C) while overexpression of dPIP_4_K resulted in lower steady-state levels of PIP_3_ (Fig. 7D). These findings are consistent with the results from our imaging of PIP_3_ levels from fixed salivary glands of these genotypes (see Fig 3 A(ii) and E). We also analyzed the rate of change in PIP_3_ levels during the initial phase following insulin stimulation. This analysis clearly revealed an enhanced rate of PIP_3_ production on loss of *dPIP_4_K*^*29*^ relative to controls and a reduced rate of PIP_3_ production in cells overexpressing this enzyme (Fig. 7E(i)). Thus, dPIP_4_K has the ability to modulate the rate of PIP_3_ production during insulin stimulation. A similar analysis of the rate of decrease in PIP_3_ levels during the phase after wortmannin addition (i.e when Class I PI_3_K activity has been inhibited) showed a marginally slower rate of decay in PIP_3_ levels in both dPIP_4_K depleted cells relative to controls but also in cells overexpressing dPIP_4_K (Fig. 7E(ii)). This finding implies that while dPIP_4_K function is able to modulate the PIP_3_ phosphatase activity operative during insulin signalling in *Drosophila* salivary gland cells, it has significantly greater effect on the Class I PI_3_K activity.

### *dPIP_4_K* function regulates sugar metabolism during larval development

We tested if increased sensitivity to insulin seen in *dPIP_4_K*^*29*^ had any impact on the physiological response of the animals to sugar intake. It has previously been reported that larvae raised on a high sugar diet (HSD) develop an insulin resistance phenotypes reminiscent of Type II diabetes [42,43]. At the level of the organism, this includes reduced body weight, a developmental delay and elevated levels of hemolymph trehalose, the main circulating sugar in insect hemolymph. As previously reported, we found that when grown on HSD (1M sucrose), wild-type larvae show ca. 9 days delay in development compared to animals grown on normal food (0.1M Sucrose) (Fig. 8A). However, interestingly, in *dPIP_4_K*^*29*^ larvae grown on HSD a delay of only 5 days was seen compared to the same genotype grown on 0.1M sucrose (Fig. 8A). We also biochemically measured the levels of circulating trehalose in the hemolymph of wandering third instar larvae. It was observed that *dPIP_4_K*^*29*^ larvae raised on normal food, showed circulating trehalose levels are *ca.* 40% lower compared to controls. Further, when wild-type animals were grown on HSD, circulating trehalose levels in larvae were elevated by ca. 25 % compared to that on normal food (Fig. 8B). However, when *dPIP_4_K*^*29*^ larvae were raised on HSD, circulating trehalose levels remained essentially unchanged (Fig. 8B) compared to that in animals grown on normal food. Together, these observations suggest that increased insulin sensitivity occurring upon loss of dPIP_4_K confers partial protection against phenotypes that arise when larvae are challenged with a high sugar diet.

**Fig. 8.**
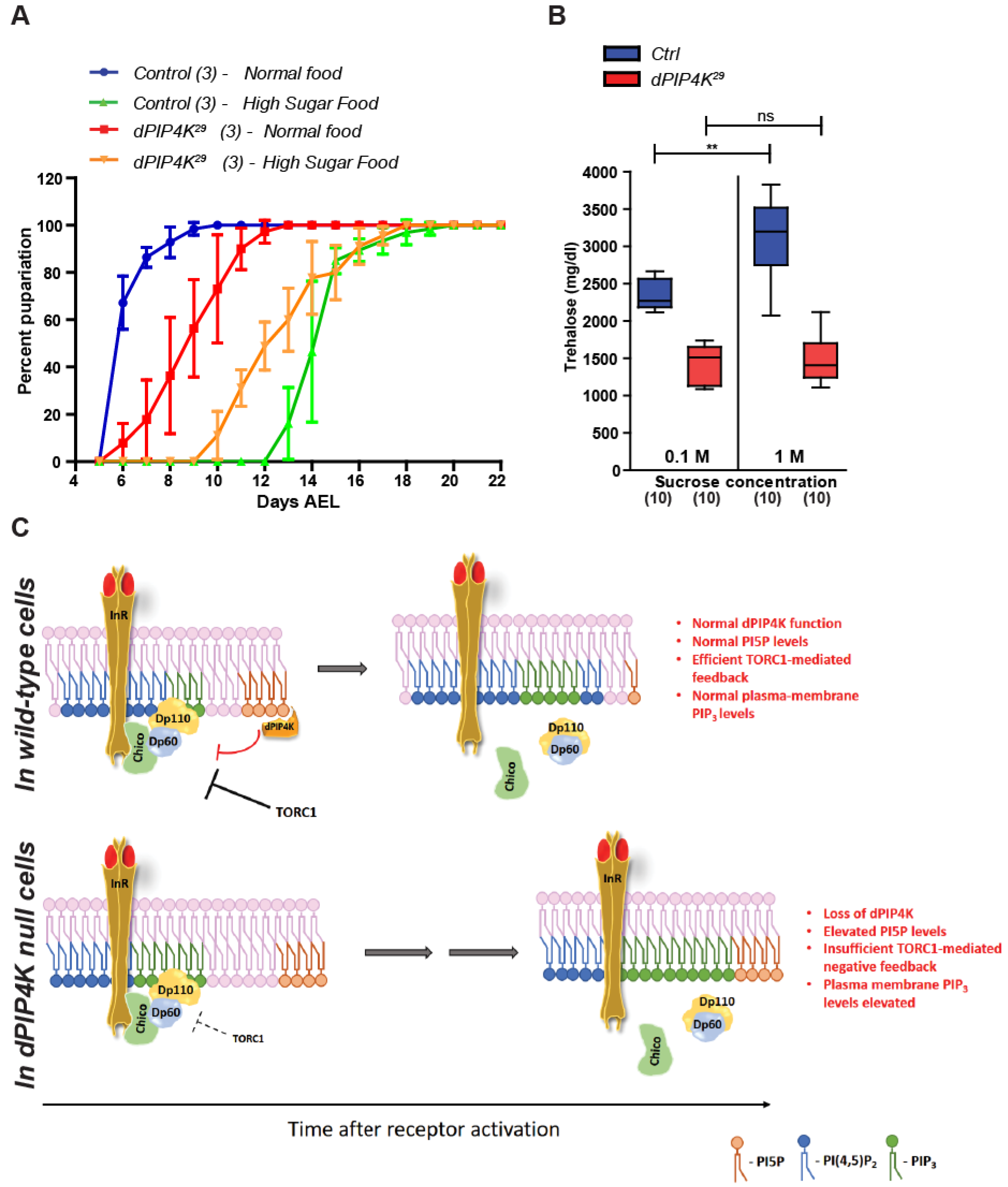
dPIP_4_K modulates the acquisition of insulin resistance upon high dietary sugar intake. **A.** The graph represents the mean percentage of pupariation (After egg laying, AEL) observed over time on indicated diets. Data collected from 3 independent batches of about 15-25 larvae per batch (for *ROR and dPIP_4_K*^*29*^). Error bars indicate SD. **B.** Mean hemolymph trehalose levels measured of hemolymph pooled from 5-8 larvae each per genotype (*ROR and dPIP_4_K*^*29*^). Whiskers in the box plots represent minimum and maximum values, with a line at the median. Numbers inside the parentheses below the plots indicate the no. of biological replicates used for the measurement. *Mann Whitney test* used for statistical analysis of the distributions. \*\**p-value* <0.01. **C.** A model for regulation of PIP_3_ levels by dPIP_4_K upon insulin stimulation. In wild-type cells, insulin-induced activation of the receptor triggers dPIP_4_K activity. This prevents PI_5_P elevation at plasma membrane and also initiates events that prevent sustained Class I PI_3_K activity and PIP_3_ production. Eventually, the negative feedback via TORC_1_ activity also sets in. These events act together and keep PIP_3_ levels in check. Upon loss of dPIP_4_K in cells, PI_5_P accumulates. Class I PI_3_K shows has a sustained activity resulting increased PIP_3_ levels upon insulin stimulation which cannot be completely restored via TORC_1_-mediated negative feedback.

## Discussion

The generation of the signalling lipid PIP_3_, is a conserved element of signal transduction by many growth factor receptors. The enzymes that control PIP_3_ levels during this process, namely Class I PI_3_K and the lipid phosphatases PTEN and SHIP2 are well studied and the biological consequences of mutations in genes encoding these enzymes underscore the importance of tight regulation of PIP_3_ levels during growth factor signalling. While the roles of many of the core enzymes that are directly involved in PIP_3_ metabolism have been studied extensively, the function of proteins that regulate their activity remains less understood; to date regulation of Class I PI_3_K activity by small GTPases (Ras, Rac) and G_βγ_ subunits has been described [reviewed in [44]]. Although a role for PIP_4_K enzymes in regulating growth factor signalling through PIP_3_ generation has been reported by earlier studies [15,16,18], the biochemical mechanism and cell-biological context in which they do so has remained obscure. PIP_4_Ks convert PI_5_P to PI(4,5)P_2_ but to date no study has found a role for PIP_4_K in regulating overall cellular PI(4,5)P_2_ levels [reviewed in [12]]. One possibility that has been raised is that PIP_4_Ks may generate the PI(4,5)P_2_ pool from which PIP_3_ is produced by Class I PI_3_K activity. Although PI_5_P, the preferred substrate of PIP_4_K, is a low abundance lipid, in principle, it is possible that a small, local pool of PI(4,5)P_2_ is generated from PI_5_P by PIP_4_K and the loss of this small pool of PI(4,5)P_2_ is not detected by the mass assays for estimating total cellular levels of this lipid. Quantitatively, based on their relative abundance, the small PIP_4_K generated pool of PI(4,5)P_2_is likely to be sufficient to serve as the substrate for PIP_3_ generation by Class I PI_3_K. A recent study [18] has reported that PIP_3_ levels are reduced in immortalized B-cells in which PIP_4_K_2_A activity is down regulated. By contrast, it has been previously reported that loss of PIP_4_K_2_B in mice results in increased levels of insulin signalling readouts such as pAKT^308^ that are direct correlates of PIP_3_ levels [15]. The exact reasons for these conflicting results is unclear and may include the different cell types used in each study; a key reason is likely to be the overlapping function of the three PIP_4_K isoforms present in mammalian genomes. In this study, we found that in *Drosophila*, that contains only a single gene encoding PIP_4_K activity (dPIP_4_K)[11,45], the levels of plasma membrane PIP_3_ in cells lacking dPIP_4_K were elevated compared to controls. We established this finding using both a fluorescent reporter for plasma membrane PIP_3_ in single cell assays using multiple cell types and also using lipid mass spectrometry across larval tissues and cultured, dPIP_4_K depleted *Drosophila* S2 cells. Thus, our study clearly demonstrates that in *Drosophila*, dPIP_4_K autonomously functions as a negative regulator of PIP_3_ production during growth factor stimulation. The elevated PIP_3_ levels seen when dPIP_4_K is depleted are not consistent with a role for this enzyme in generating the PI(4,5)P_2_ at the plasma membrane used by Class I PI_3_K as substrate to generate PIP_3_ during insulin signalling. Therefore, is likely that dPIP_4_K regulates PIP_3_ levels through its ability to control the function of proteins that themselves regulate PIP_3_ levels during Class I PI_3_K signalling.

In an earlier study [11], we had observed (*dPIP_4_K*^*29*^ larvae have systemically reduced mTOR activation. In mammalian cells, reducing mTOR activity through the use of rapamycin or a loss of S6K, a direct target of TORC_1_, leads to increased activation of insulin signalling and obesity resistance associated with increased insulin sensitivity [46–48]. It is also reported that S6K inactivates IRS-1 by phosphorylating it on multiple serine residues [28,36]. Therefore, it is reasonable to hypothesize a scenario where the reduced TORC_1_ activity in *dPIP_4_K*^*29*^ cells may be the defect that drives the increase in the levels of PIP_3_ in *dPIP_4_K*^*29*^. In wild-type larval cells, modulating TORC_1_ activity had expected effects on cell size (Fig. S5A(i), B(i) and C(i)) but also could tune PIP_3_ levels during insulin stimulation (Fig. 5A, C, E); enhancing TORC_1_ output resulted in lower levels of insulin-induced PIP_3_ whereas reducing TORC_1_ activity caused higher levels of PIP_3_. By contrast, overexpression of Rheb or the down-regulation of Tsc1/2 was not able to revert the elevated plasma membrane PIP_3_ levels in *dPIP_4_K*^*29*^ cells (Fig. 5B, D and F) although they were able to restore the reduced cell size in *dPIP_4_K*^*29*^ (Fig. S5A(ii) and S5C(ii)). These results imply two conclusions: 1) Decreased TORC_1_ activity is not sufficient to explain the enhanced PIP_3_ levels in *dPIP_4_K*^*29*^ larval cells and 2) Efficient feedback regulation of PIP_3_ levels during receptor tyrosine kinase activation requires intact dPIP_4_K function in addition to TORC_1_ activity.

Binding of insulin to its receptor triggers a signalling cascade where the initial events occur at the plasma membrane. These involve interaction of the activated insulin receptor-ligand complex with IRS followed by the recruitment and activation of Class I PI_3_K at the plasma membrane. At which sub-cellular location is dPIP_4_K activity required to regulate this process? Fractionation and immunolocalization studies in mammalian cells [40] and *Drosophila* [11] have indicated that PIP_4_K isoforms are distributed across multiple subcellular compartments including the plasma membrane, nucleus and internal vesicular compartments. In this study, using selective reconstitution of the dPIP_4_K to specific membrane compartments, in cells devoid of any endogenous PIP_4_K protein, we found that plasma membrane targeted dPIP_4_K could rescue the elevated PIP_3_ levels in dPIP_4_K null cells. This observation strongly suggests that the plasma membrane localized dPIP_4_K is sufficient to control PIP_3_ production during insulin stimulation. Our observation that *dPIP_4_K*^*29*^ cells were hypersensitive to overexpression of *dINR* or *chico* compared to wild-type cells likely reflects the loss of a dPIP_4_K dependent event in the control of PIP_3_ levels at the plasma membrane. Overexpression of plasma-membrane localized PIP_4_K_2_B was able to reduce pAKT^308^ phosphorylation upon insulin stimulation in mammalian cells just as well as the wildtype PIP_4_K_2_B enzyme. Our finding of a role for the plasma membrane localized PIP_4_K in regulating PIP_3_ levels in both *Drosophila* and mammalian cells underscores the evolutionarily conserved nature of this mechanism. Previous studies have shown that levels of PI_5_P, the substrate for PIP_4_KS, increases upon insulin stimulation and importantly, addition of exogenous PI_5_P can stimulate glucose uptake in a PI_3_K-dependent manner [17,49]. Therefore, plasma membrane localized PIP_4_K and the levels of its substrate PI_5_P could be a mechanism by which early events during insulin signalling are regulated.

What molecular event involved in PIP_3_ turnover might dPIP_4_K regulate at the plasma membrane? Using live cell imaging studies of PIP_3_ turnover at the plasma membrane coupled with chemical inhibition of Class I PI_3_K, we were able to observe that dPIP_4_K function has a substantial impact on the rate of PIP_3_ production following insulin stimulation whereas the rate of PIP_3_ degradation was only marginally affected. This finding suggests that dPIP_4_K likely regulates the activity of Class I PI_3_K either directly or by controlling its coupling to the activated insulin receptor complex at the plasma membrane; the exact mechanism by which it does so remains to be established.

What might be the physiological consequence of losing dPIP_4_K mediated feedback control on PIP_3_ production in the context of insulin signalling? Previous studies in mouse and human cells have reported that excessive activation of TORC_1_ signalling leads to inactivation of insulin signalling pathway and development of insulin resistance [50–52]. Since TORC_1_ activity is reduced [11] and PIP_3_ were elevated (this study) in animals lacking dPIP_4_K, it is likely that loss of dPIP_4_K impacts sugar metabolism in *Drosophila* larvae. Using a recently reported high-sugar induced obesity and Type II diabetes-like disease model in *Drosophila* [42], we found that *dPIP_4_K*^*29*^ larvae appear resistant to a high sugar diet as measured by the elevation in the hemolymph trehalose levels and they were relatively resistant to the developmental delay seen when wild-type larvae are reared on a high-sugar diet. This observation is reminiscent of that reported for the PIP_4_K_2_B^−/−^ mice that have a reduced adult body weight compared to controls and clear blood glucose faster following a sugar bolus than control animals [15]. Our observation that dPIP_4_K at the plasma-membrane controls sensitivity to insulin receptor activation suggests a molecular basis for the physiological phenotypes observed in *dPIP_4_K*^*29*^ larvae and PIP_4_K_2_B^−/−^ mice. These observations also raise the possibility that inhibition of PIP_4_K activity may offer a route to reducing insulin resistance in the context of Type II diabetes. Such a mechanism may explain the hyperactivation of the T-cell receptor responses in mice lacking PIP_4_K_2_C [14], since the activation of Class I PI_3_K is a key element of T-cell receptor signal transduction. More generally, PIP_4_K activity likely offers a novel element of regulation for Class I PI_3_K activity in the context of receptor tyrosine kinase signalling.

## Materials and Methods

### *Drosophila* strains and rearing

Unless indicated, flies were grown on standard fly medium containing corn meal, yeast extract, sucrose, glucose, agar and antifungal agents. For all experiments, crosses were setup at 25°C in vials/bottles under non-crowded conditions.

Fly medium composition:

| Ingredients | 0.1 M sucrose | 1 M sucrose |
| --- | --- | --- |
| For 1 Litre |  |  |
| Corn flour | 80 g | 80 g |
| D-Glucose | 20 g | 20 g |
| Sugar | 40 g | 342 g |
| Agar | 8 g(4g) | 8 g |
| Yeast powder | 15 g | 15 g |
| Propionic acid | 4 ml | 4 ml |
| TEGO(Methyl parahydroxy benzoate) | 0.7 g | 0.7 g |
| Orthophosphoric acid | 0.6 ml | 0.6 ml |

The following stocks were used in the study: wildtype strain *Red Oregon R (ROR), AB1-Gal_4_* (Bloomington # 1824), *UAS-dInR* (Bloomington # 8262), *UAS-Rheb*^*RNAi*^ (Bloomington TRiP # 33966), *UAS-Rheb* (Bloomington # 9688), UAS-TSC^RNAi^(Bloomington TRiP # 52931), P{tGPH}4 (Bloomington # 8164), UAS-*dPIP_4_K*^*RNAi*^ (Bloomington TRiP # 65891). *UAS-dPIP_4_K::eGFP* and *dPIP_4_K*^*29*^ were generated in the lab and described in [11]. For PIP_3_ measurements in the *dPIP_4_K*^*29*^ rescue experiment (Fig. 4F) using GFP-PH-GRP1 probe, we cloned dPIP_4_K cDNA (BDGP clone#LD10864) into pUAST-attB between *EcoRI* and *Xhol* sites without the GFP tag. The generation of flies expressing dPIP_4_K::-mCherry-CAAX is described in *Kumari K et.al, 2017*. For targeting dPIP_4_K to the endomembranes, the sequence QGSMGLPCVVM (*Sato.M et. al, 2006*) replaced the CAAX motif in the dPIP_4_K::mCherry-CAAX construct. To generate Lysosomal-dPIP_4_K::eGFP, the 39 amino-acid sequence from p18/LAMTOR (*Menon S et.al., 2014*) was used as a signal sequence. The signal sequence was commercially synthesized with a C-terminal flag tag and introduced upstream of dPIP_4_K::eGFP. The entire fusion construct was cloned into pUAST-attB by GIBSON assembly using *NotI* and *XbaI* sites. All molecular constructs conceptualized and analysed further with use of the molecular cloning tools available on the free online platform ‒ Benchling.com. All transgenic lines were generated using insertions that were performed using site-specific recombination. The level of GFP fluorescence from lysosomal-dPIP_4_K::eGFP was observed to be very low in the salivary glands and did not interfere with our analysis PIP_3_ measurements using the GFP-PH-GRP1 probe in Fig. 6F. All primer sequences used for cloning the different constructs are available on request.

### Cell Culture, dsRNA treatment and Insulin stimulation assays

CHO cell line stably expressing insulin receptor (isoform A) was a kind gift from Dr Nicholas Webster, UCSD. These were maintained at standard conditions in HF12 culture medium supplemented with 10% Fetal bovine serum and under G418 selection (400 μg/ml). Transfections were done 48 hrs. before the assay using FuGene, Promega Inc. as per manufacturer’s protocols when the cultures were 50% confluent. For insulin stimulation assays, cells were starved overnight in HF12 medium without serum. Thereafter, the cells were de-adhered, collected into eppendorf tubes and stimulated with 1 μM insulin for indicated times. Post stimulation, cells were spun down and immediately lysed.

S2R+ cells were cultured in Schneider’s medium (GIBCO 21720024, HiMedia Labs IML003A) supplemented with 10% non-heat inactivated fetal bovine serum (US Origin, GIBCO 16000044) and contained antibiotics ‒ streptomycin and penicillin (SIGMA G1146). dsRNA was synthesised inhouse using Megascript RNAi Kit (Ambion, Life Technologies, AM1626) as per manufacturer’s instructions. For dsRNA treatments, 0.5 × 10^6^ cells were seeded into a 24-well plate. Once observed to be settled, cells were incubated with incomplete medium containing 1.875 μg of dsRNA. After 1 hour, an equal amount of complete medium was added to each well. The same procedure was repeated on each well 48 hours after initial transfection after removal of the spent medium from each well. Cells were harvested by trypsinization after a total of 96 hours of dsRNA treatment. For mass spectrometric estimation of PIP_3_, S2R+ cells were pelleted down and stimulated with 1 μM insulin for 10 min. The reaction was stopped by the addition of ice-cold initial organic mix (described later in the section) and used for lipid extraction. The primer sequences used for dsRNA preparation are-

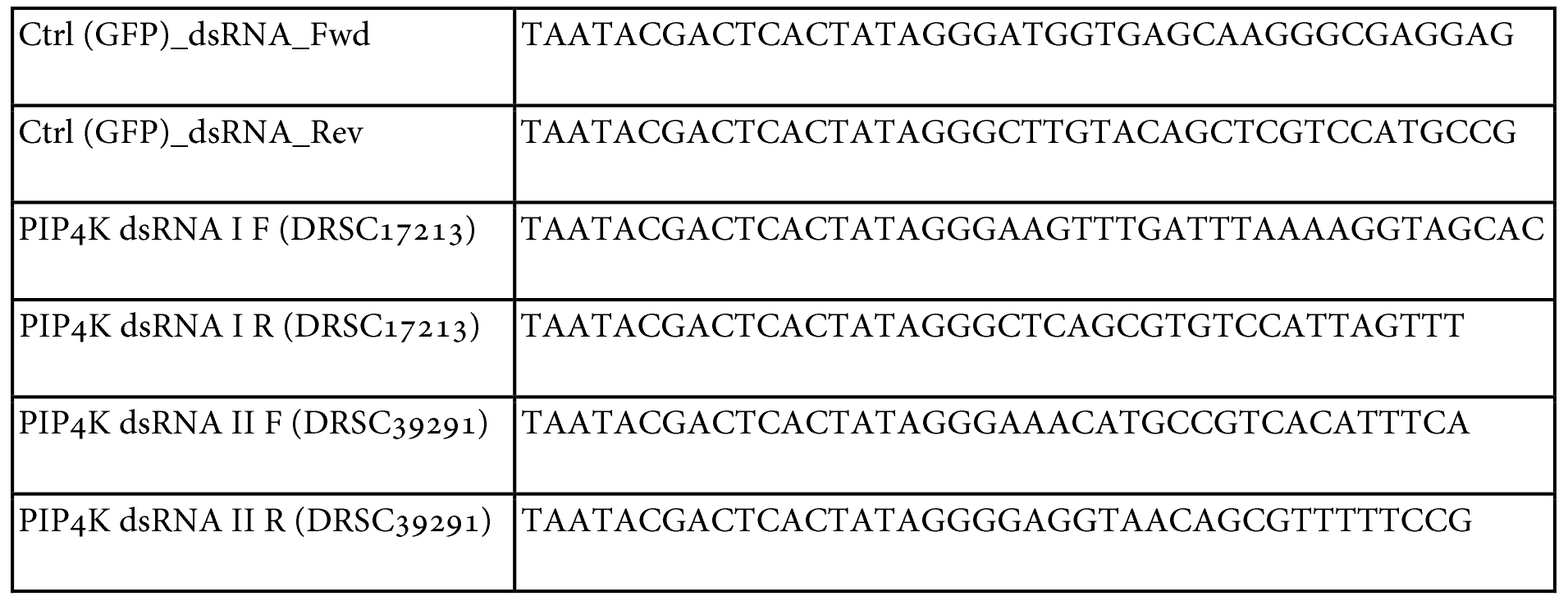

### Larval growth Curve Analysis

Adult flies were made to lay eggs within a span of 4-6 hrs on normal food. After 24 hrs, newly hatched first instar larvae were collected and transferred in batches of about 15-25 larvae per into vials containing either 0.1/1M Sucrose in the fly media with other components unaltered. The vials were then observed to count the number of pupae.

### Hemolymph Trehalose Measurements

The measurements were done exactly as described in [42]. In brief, hemolymph was pooled from five to eight larvae to obtain 1 μl for assay. The reagents used porcine trehalase (SIGMA, T8778) and GO kit (SIGMA, GAGO20)

### Cell size analysis in salivary glands

Salivary glands were dissected from wandering third instar larvae and fixed in 4% paraformaldehyde for 30 min at 4°C. Post fixation, glands were washed thrice with 1X PBS and incubated in BODIPY-FL-488 for 3 hours at room temperature. The glands were washed thrice in 1X PBS following which nuclei were labelled (using either DAPI or TOTO_3_) for 10 mins at room temperature and washed with 1X PBS again. The glands were then mounted in 70% glycerol and imaged within a day of mounting. Imaging was done on Olympus FV1000 Confocal LSM using a 20X objective. The images were then stitched into a 3D projection using an ImageJ plugin. These reconstituted 3D z-stacks were then analysed for nuclei numbers (correlate for cell number) and volume of the whole gland using Volocity Software (version 5.5.1, Perkin Elmer Inc.). The average cell size was calculated as the ratio of the average volume of the gland to the number of nuclei.

### *Ex vivo* insulin stimulation and PIP_3_ measurements in salivary glands and fat body

For experiments with salivary glands, wandering third instar larvae were dissected one larva at a time and glands were immediately dropped into a well of a 96-well plate containing either only PBS or PBS + 10 μM Insulin (75 μl) and incubated for 10 min at RT. Following this, 25 μl of 16% PFA was added into the same well to yield a final conc. of 4% PFA. The glands were fixed in this solution for 18 min at room temperature and then transferred sequentially to wells containing PBS every 10 min for 3 washes. Finally, glands were mounted in 80% glycerol in PBS containing antifade (0.4% propyl-gallate). For experiments with fat body lobes, late third instar feeding larvae were starved by placing them on a filter paper soaked in lX PBS for 2 hrs. Thereafter, the incubation, fixation and mounting steps were done exactly as described for salivary glands. Imaging was done on LSM 780 inverted confocal microscope with a 20X/0.8 NA Plan Apochromat objective. For quantification, confocal slices were manually curated to generate maximum z-projections of middle few planes of cells. Thereafter, line profiles were drawn across clearly identifiable plasma membrane regions and their adjacent cytosolic regions and ratios of mean intensities for these line profiles were calculated for each cell. For salivary glands, about 10-15 cells from multiple glands were analysed and used to generate statistics. For fat body, about 50 cells each from multiple animals were used for analysis.

For live imaging, salivary glands from wandering third instar larvae were dissected (glands from one larva imaged in one imaging run) and placed inside a drop of imaging buffer (lX PBS containing 2mg/ml glucose) on a coverglass. The buffer was carefully and slowly soaked out with a paper tissue to let the glands settle and adhere to the surface. Thereafter, the glands were immediately rehydrated with 25 μl of imaging buffer. The imaging was done on Olympus FV3000 LSM confocal system using a 10X objective. A total of 80 frames of a single plane were acquired, with 10s intervals. While imaging, 25 μl of 20uM (2X) bovine insulin was used to stimulate the glands. After the steady state was achieved, 50 μl of 8oonM (2X) of wortmannin was added on top to inhibit PI_3_K activity.

### PIP_3_ measurement by LC-MS/MS

The method was adopted and modified as required from [34].

### Lipid extraction

5 larvae were dissected in 1X PBS and transferred immediately into 37.5 μl of 1X PBS in a 2 ml Eppendorf. For insulin stimulation, to this, 37.5 μl of 100 μM Insulin (final concentration ‒ 50 μM) was added and the tube was incubated on a mix mate shaker for 10 min at 500 rpm. At the end of incubation time, 750 μl of ice-cold 2:1 MeOH:CHCl_3_ organic mix was added to stop the reaction. Part of this solution was decanted and the rest of the mix containing larval tissues was transferred into a homogenization tube. Larval tissues were homogenized in 4 cycles of 10 secs with 30 sec intervals at 6000 rpm in a homogenizer (Precellys, Bertin Technologies). The tubes were kept on ice at all intervals. The entire homogenate was then transferred to a fresh eppendorf and the homogenization tube was then washed with the decanted mix kept aside earlier. 120 μl of water was added to the homogenate collected in eppendorf, followed by addition of 5 ng of 17:0, 20:4 PIP_3_ internal standard (ISD). The mixture was vortexed and 725 μl of chloroform was added to it. After vortexing again for 2 min at around 1000-1500 rpm, the phases were separated by centrifugation for 3 min at 1500g. 1ml of lower organic phase was removed and stored in a fresh tube. To the remaining aqueous upper phase, again 725 μl of chloroform was added. The mixture was vortexed and spun down to separate the phases. Again, 1 ml of the organic phase was collected and pooled with the previous collection (total of 2ml). This organic phase was used for measuring total organic phosphate. To the aqueous phase, 500 μl of the initial organic mix was added followed by 170 μl of 2.4M HC1 and 500 μl of CHC1_3_. This mixture was vortexed for 5 min at 1000-1500 rpm and allowed to stand at room temperature for 5 minutes. The phases were separated by centrifugation (1500g, 3 min). The lower organic phase was collected into a fresh tube by piercing through the protein band sitting at the interface. To this, 708 μl of lower phase wash solution was added, the mixture was vortexed and spun down (1500g, 3 min). The resultant lower organic phase was completely taken out carefully into an Eppendorf tube and used for derivatization reaction.

### Extraction solvent mixtures

Initial organic mix: MeOH/Chloroform in the ratio of 484/242 ml, Lower Phase Wash Solution: Methanol/1 M hydrochloric acid/ chloroform in a ratio of 235/245/15 ml. All ratios are expressed as vol/vol/vol.

### Derivatization of Lipids

To the organic phase of the sample, 50 μl of 2M TMS-Diazomethane was added (TO BE USED WITH ALL SAFETY PRECAUTIONS!). The reaction was allowed to proceed at room temperature for 10 min at 600 rpm. After 10 min, 10 μl of Glacial acetic acid was added to quench the reaction, vortexed briefly and spun down. 700 μl of post derivatization wash solvent was then added to the sample, vortexed (2 min, 1000-1500 rpm) and spun down. The upper aqueous phase was discarded and the wash step was repeated. To the final organic phase, 100 μl of 9:1 MeOH:H_2_O mix was added and the sample was dried down to about 10-15 μl in a speedvac under vacuum.

### Chromatographic separation and Mass spectrometric detection

The larval lipid extracts were re-suspended in 170 μl LC-MS grade methanol and 30 μl LC-MS grade water. Samples were injected as duplicate runs of 3.5 μl. Chromatographic separation was performed on an Acquity UPLC BEH300 C4 column (100 × 1.0 mm; 1.7 μm particle size) purchased from Waters Corporation, USA on a Waters Aquity UPLC system and detected using an ABSCIEX 6500 QTRAP mass spectrometer. The flow rate was 100 μL/min. Gradients were run starting from 55% Buffer A (Water + 0.1% Formic Acid)- 45% Buffer B (Acetonitrile + 0.1% Formic acid) to 42% B from 0-5 min; thereafter 45% B to 100% B from 5-10 min; 100% B was held from 10-15 min; brought down from 100% B to 45% B between 15-16 min and held there till 20th min to re-equilibrate the column. On the mass spectrometer, in pilot standardization experiments, we first employed Neutral Loss Scans on biological samples to look for parent ions that would lose neutral fragments of 598 a.m.u indicative of PIP_3_ lipid species (as described in [34]). Thereafter, these PIP_3_ species were quantified in biological samples using the selective Multiple Reaction Monitoring (MRM) method in the positive mode. Only those MRM transitions that showed an increase upon insulin stimulation of biological samples were used for the final experiments (depicted in figure S3B). The MRM transitions for the different PIP_3_ species quantified are listed out in the table below. Area of all the peaks was calculated on Sciex MultiQuant software. The area of the internal standard peak was used to normalize for lipid recovery during extraction. The normalized for each of the species was then divided by the amount of organic phosphate measured in each of the biological samples. The other mass spectrometer parameters are as follows: ESI voltage: +4500V; Dwell time: 40 ms; DP (Declustering Potential): 35.0 V; EP: (Entrance Potential): 10.1 V, CE (Collision Energy): 47.0 V; CXP (Collision cell Exit Potential): 11.6 V, Source Temperature : 450 C, Ion Spray Voltage ‒ 4000 V, Curtain Gas : 35.0, GS1: 15, GS2: 16. The area under the peaks was extracted using MultiQuant vi.i software (ABSCIEX). Numerical analysis was done in Microsoft Excel.

|  | Sample |  |  |  |  |  |
| --- | --- | --- | --- | --- | --- | --- |
|  | Drosophila Larvae |  |  | S2R+ cells |  |  |
|  | PIP<sub>3</sub> species | Parent Ion | Daughter Ion | PIP<sub>3</sub> species | Parent Ion | Daughter Ion |
| MRM Transitions | 32_1 | 1145.5 | 547.5 | 30_1 | 1119.5 | 521.5 |
|  | 32_2 | 1177.5 | 579.5 | 32_0 | 1149.5 | 551.5 |
|  | 34_1 | 1175.5 | 577.5 | 32_1 | 1147.5 | 549.5 |
|  | 34_2 | 1173.5 | 575.5 | 32_2 | 1145.5 | 547.5 |
|  | 34_3 | 1171.5 | 573.5 | 34_0 | 1177.5 | 579.5 |
|  | 36_1 | 1203.5 | 605.5 | 34_1 | 1175.5 | 577.5 |
|  | 36_2 | 1201.5 | 603.5 | 34_2 | 1173.5 | 575.5 |
|  | 36_3 | 1199.5 | 601.5 | 34_3 | 1171.5 | 573.5 |
|  | 36_4 | 1197.5 | 599.5 | 34_4 | 1169.5 | 571.5 |
|  |  |  |  | 34_5 | 1167.5 | 569.5 |
|  |  |  |  | 36_2 | 1201.5 | 603.5 |
|  |  |  |  | 36_3 | 1199.5 | 601.5 |
|  |  |  |  | 36_4 | 1197.5 | 599.5 |
|  |  |  |  | 36_5 | 1195.5 | 597.5 |
|  |  |  |  | 38_3 | 1227.5 | 629.5 |
|  |  |  |  | 38_4 | 1225.5 | 627.5 |
|  |  |  |  | 38_5 | 1223.5 | 625.5 |
|  |  |  |  | 37_4 ISD | 1211.5 | 613.5 |

### Total Organic Phosphate measurement

1 ml of the organic phase from each sample was taken into phosphate-free tubes and dried completely at 90°C. The remaining steps were performed as described in *Thakur et.al, 2016*.

### Preparation of S2R+ cell lysate for *in vitro* PI_5_P 4- kinase assay

The S2R+ cells were pelleted at looog for 10 min and washed with ice-cold PBS Twice. Cells were thereafter homogenized in lysis buffer containing 5omM Tris-Cl, pH ‒ 7.5, imM EDTA, 1mM EGTA, 1% Triton-X-100, 5omM NaF, 0.27 M Sucrose, 0.1% (β- Mercaptoethanol and freshly added protease and phosphatase inhibitors (Roche). The lysate was then centrifuged at looog for 15 min at 4 °C. Protein estimation was performed using the Bradford reagent according to the manufacturer’s instructions.

### PI_5_P4-kinase Assay

Vacuum-dried substrate lipid (6 μM PI_5_P) and 20 μM of phosphatidylserine were resuspended in 10 mM Tris pH 7.4 and micelles were formed by sonication for 2 min in a bath-sonicator. 50 μl of 2X PIPkinase reaction buffer (100 mM Tris pH 7.4, 20 mM MgCl_2_, 140 mM KC1, and 2 mM EGTA) containing 20 μM ATP, 5 [μCi [γ-^32^P] ATP and cell lysates containing ~10 μg total protein was added to the micelles. The reaction mixture was incubated at 30 °C for 16 h. Lipids were extracted and resolved by one dimensional TLC (45:35:8:2 chloroform: methanol: water: 25% ammonia). The resolved lipids were imaged using phosphorlmager.

### Western Blotting

For larval western blots, lysates were prepared by homogenizing 3 wandering third instar larvae or 5 pairs of salivary glands from third instar larvae. In the case of CHO-IR cells, pelleted cells were lysed by repeated pipetting in lysis buffer (same as described above). Thereafter, the samples were heated at 95°C with Laemli loading buffer for 5 min and loaded onto an SDS- Polyacrylamide gel. The proteins were subsequently transferred onto a nitrocellulose membrane and incubated with indicated antibodies overnight at 4°C (for actin/tubulin incubation was done at room temperature for 3 hrs.). Primary antibody concentrations used were ‒ anti- α-actin (SIGMA A5060) 1:1000; anti-dPIP_4_K 1:1000, anti ‒ GAPDH (Novus Biologicals, #IM-5143A), anti-PIP_4_KB (Cell Signaling, #9694), anti ‒ PAKTT308 (Cell Signaling, #9275), anti-AKT (Cell Signaling, # 9272). The blots were then washed thrice with Tris Buffer Saline containing 0.1% Tween-20 (0.1% TBS-T) and incubated with 1:10000 concentration of appropriate HRP-conjugated secondary antibodies (Jackson Laboratories, Inc.) for 1.5 hrs. After three washes with 0.1% TBS-T, blots were developed using Clarity Western ECL substrate on a GE ImageQuant LAS 4000 system.

## Acknowledgements

This work was supported by the National Centre for Biological Sciences, TIFR, Department of Biotechnology, Ministry of Science and Technology (India); and a Wellcome Trust-DBT India Alliance Senior Fellowship to PR. S.S is a recipient of the S.P Mukherji Fellowship from CSIR and S.M an ICMR Fellowship. We thank the NCBS *Drosophila*, Imaging and Lipidomics Facility for support.

## Supplementary Figure Legends

**Fig. S1.**
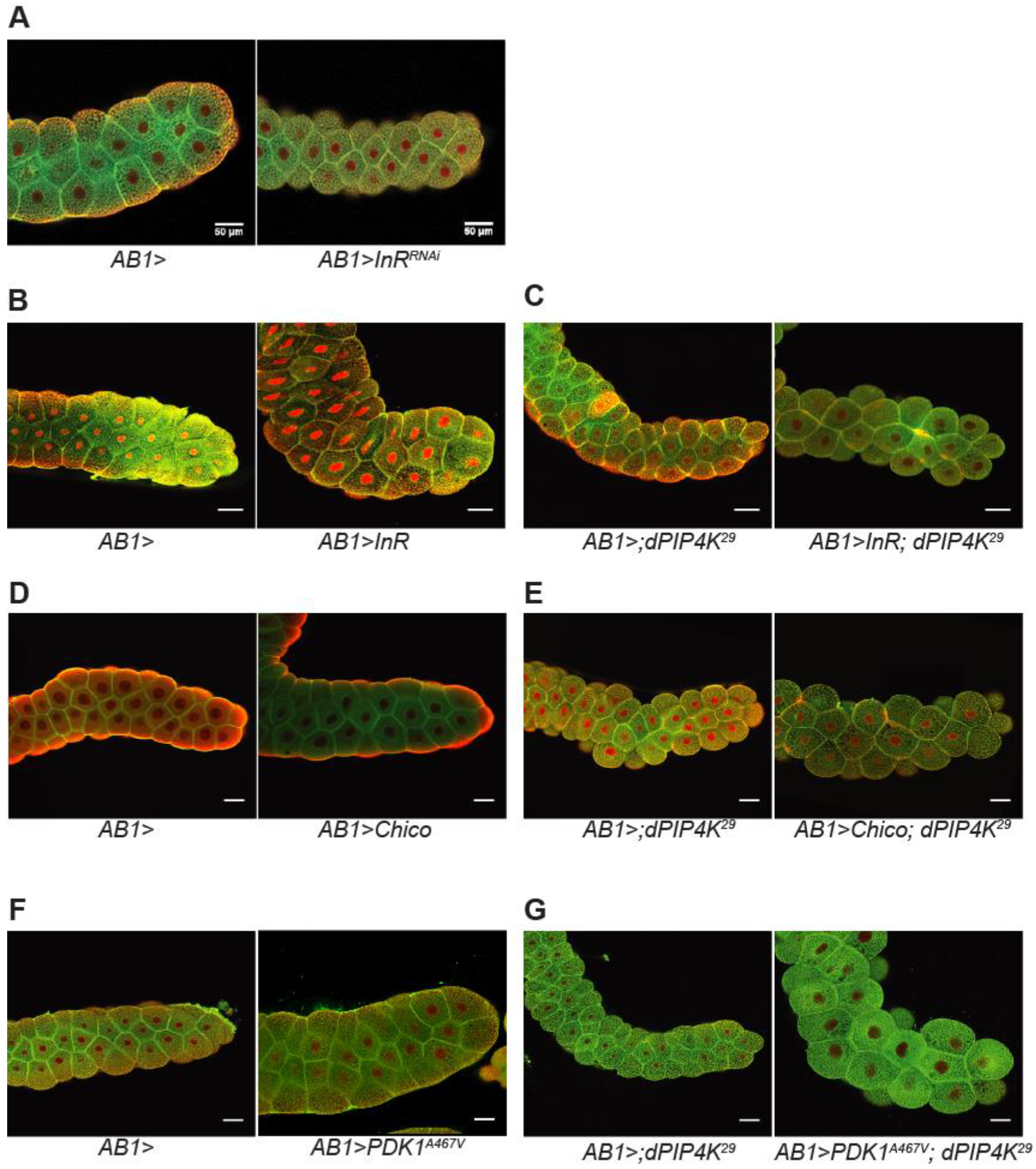
Modulatory effect of dPIP_4_K on insulin signalling in regulation of cell size. **A-G.** Representative confocal sections of salivary glands of wandering third instar larvae labelled with BODIPY FL488 (green) and TOTO3 (red) to mark the nuclei. Scale: 50 μm. Genotypes: **A.** *AB1Gal_4_*/+ *and AB1Gal_4_*/+; *UAS-dInR*^*RNAi*^. **B.** *AB1Gal_4_*/+ *and UAS-dInR*/+; *AB1Gal_4_*/+. **C.** *AB1Gal_4_*/+; *dPIP_4_K*^*29*^ *and UAS-dInR*/+; *AB1Gal_4_*/+; *dPIP_4_K*^*29*^. **D.** *AB1Gal_4_*/+ *and UAS-Chico*/+; *AB1Gal_4_*/+. **E.** *AB1Gal_4_*/+; *dPIP_4_K*^*29*^ *and UAS-Chico*/+; *AB1Gal_4_*/+; *dPIP_4_K*^*29*^. **F.** *AB1Gal_4_*/+ *and UAS-PDK*^*A467v*^;; *AB1Gal_4_*/+. **G.** *AB1Gal_4_*+; *dPIP_4_K*^*29*^ *and UAS-PDK*^*A467V*^; *AB1Gal_4_; dPIP_4_K*^*29*^.

**Fig. S2.**
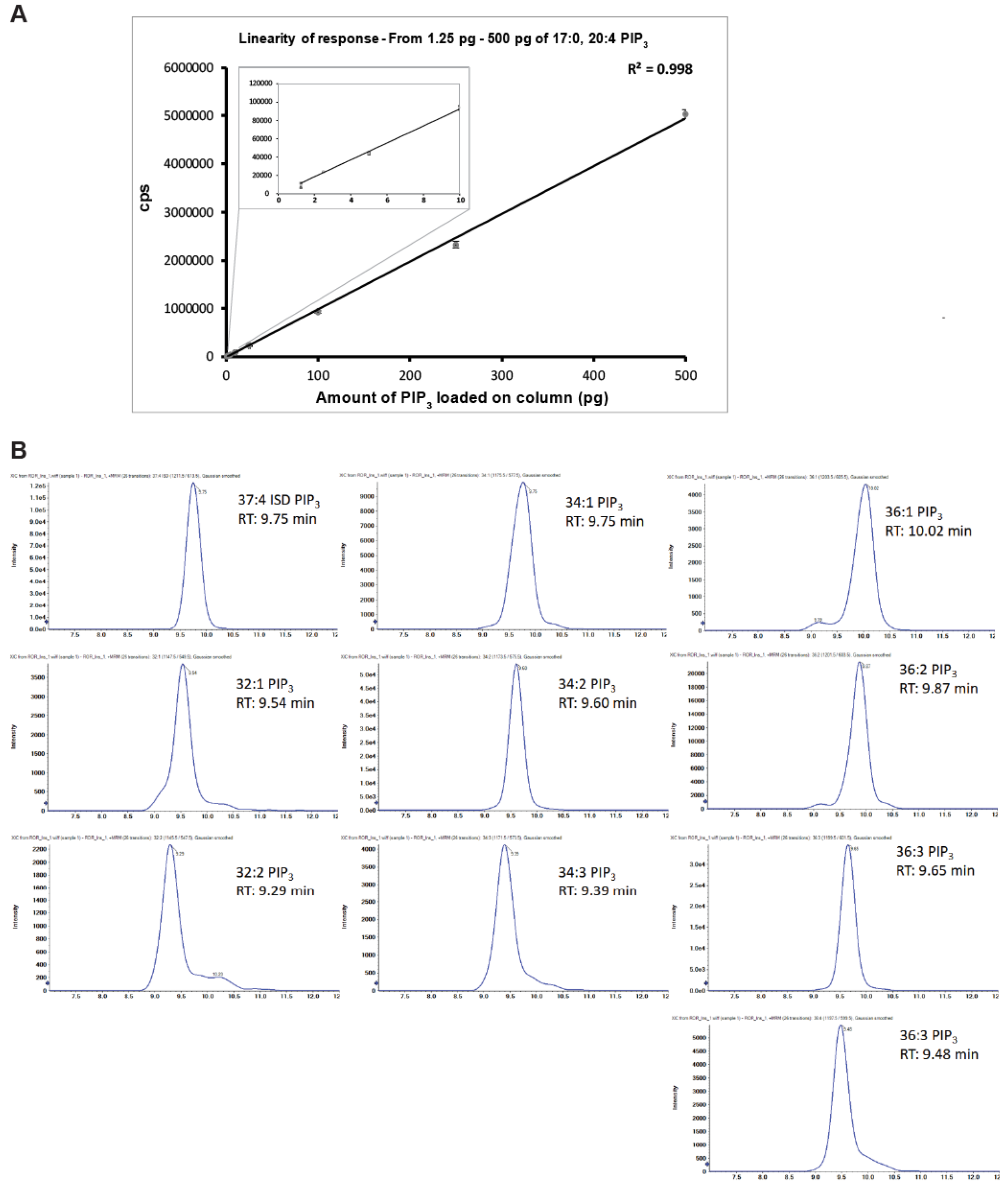
Standardisation of PIP_3_measurement using LCMS. **A.** Linearity of mass spectrometer response for increasing amounts of PIP_3_ standard (17:0, 20:4) injected. Each point on the curve indicates the mean ± SD of three replicate injections. **B.** Chromatograms showing the elution profiles and retention times for various PIP_3_ species detected from whole larval lipid extracts of wildtype larvae stimulated *ex-vivo* with 100 μM insulin for 10 min. Note the changing retention times with increase in no. of double bonds and increase in length of acyl chains. Increase in double bonds for a fixed acyl chain length results in earlier elution. Increase in length of acyl chain delays elution.

**Fig. S3.**
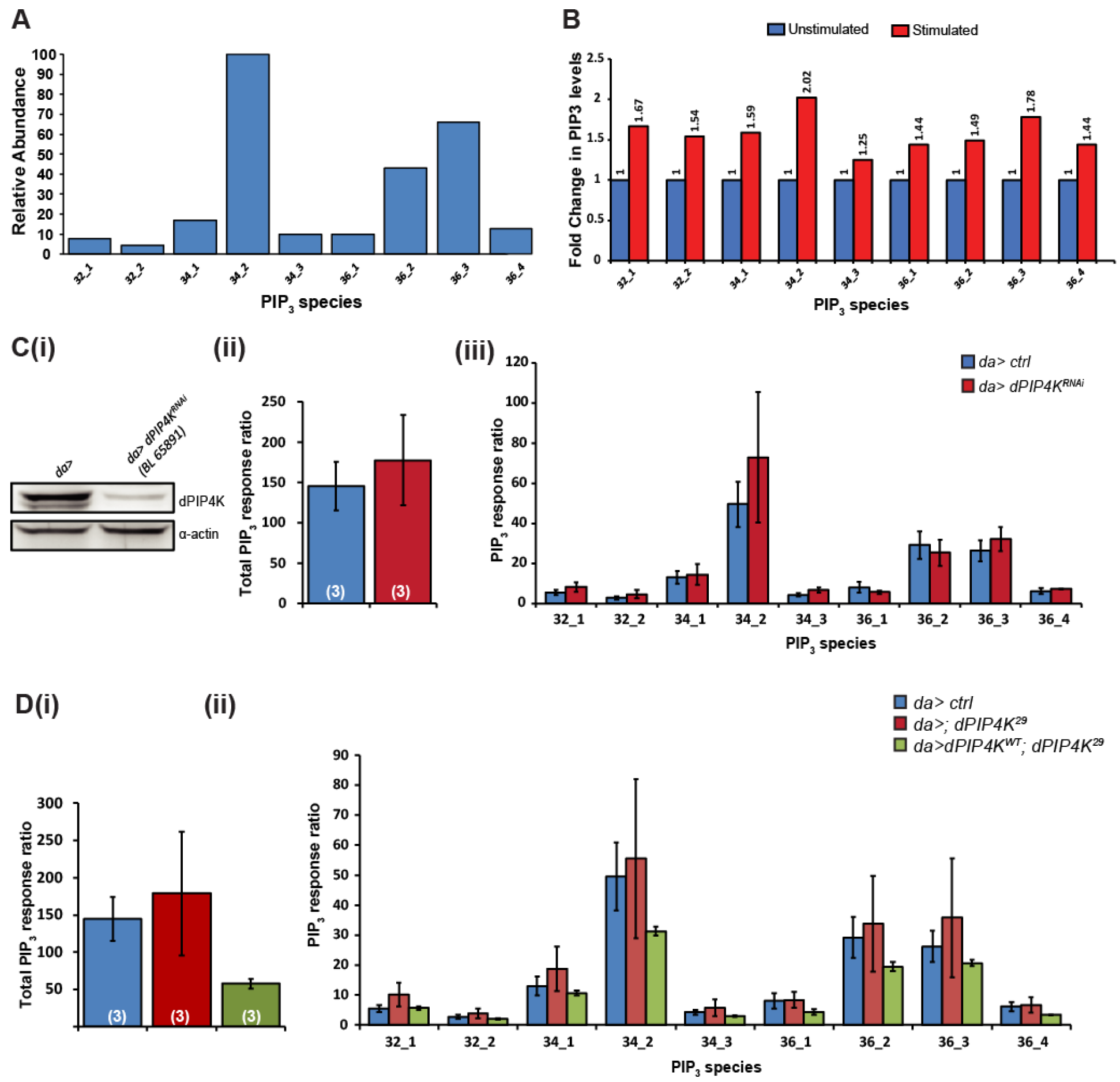
Biochemical measurement of PIP_3_ from larval extracts using LCMS. **A.** Relative abundance of various PIP_3_ species in whole larval lipid extracts of wildtype larvae stimulated *ex-vivo* with 100 μM insulin for 10 min. **B.** An experiment showing changes in the levels of various PIP_3_ species upon insulin stimulation (100 μM, 10 min). **C(i).** Immunoblot from whole larval lysates showing reduction in levels of dPIP_4_K protein upon pan-larval RNAi for dPIP_4_K (*UAS-dPIP_4_K*^*RNAi*^/+; *daGal_4_*/+). **C(ii).** Total levels of PIP_3_in whole larval control and *dPIP_4_K*^*RNAi*^ lipid extracts **C(iii).** Levels of various larval PIP_3_ species in whole larval control and *dPIP_4_K*^*RNAi*^ lipid extracts. Total PIP_3_ levels **(D(i))** and levels of individual species **(D(ii))** in Gal_4_-control (*daGal_4_*/+), Gal_4_-control in *dPIP_4_K*^*29*^ background (*daGal_4_*/+; *dPIP_4_K*^*29*^) and pan-larval rescue (*daGal_4_/UAS-dPIP_4_K::eGFP; dPIP_4_K*^*29*^) lipid extracts. The graphs show mean PIP_3_ levels (normalized to spiked internal standards and total lipid phosphates recovered). Error bars depict SD. On each panel, numbers inside the parentheses indicate the no. of biological replicates used for the measurement.

**Fig. S4.**
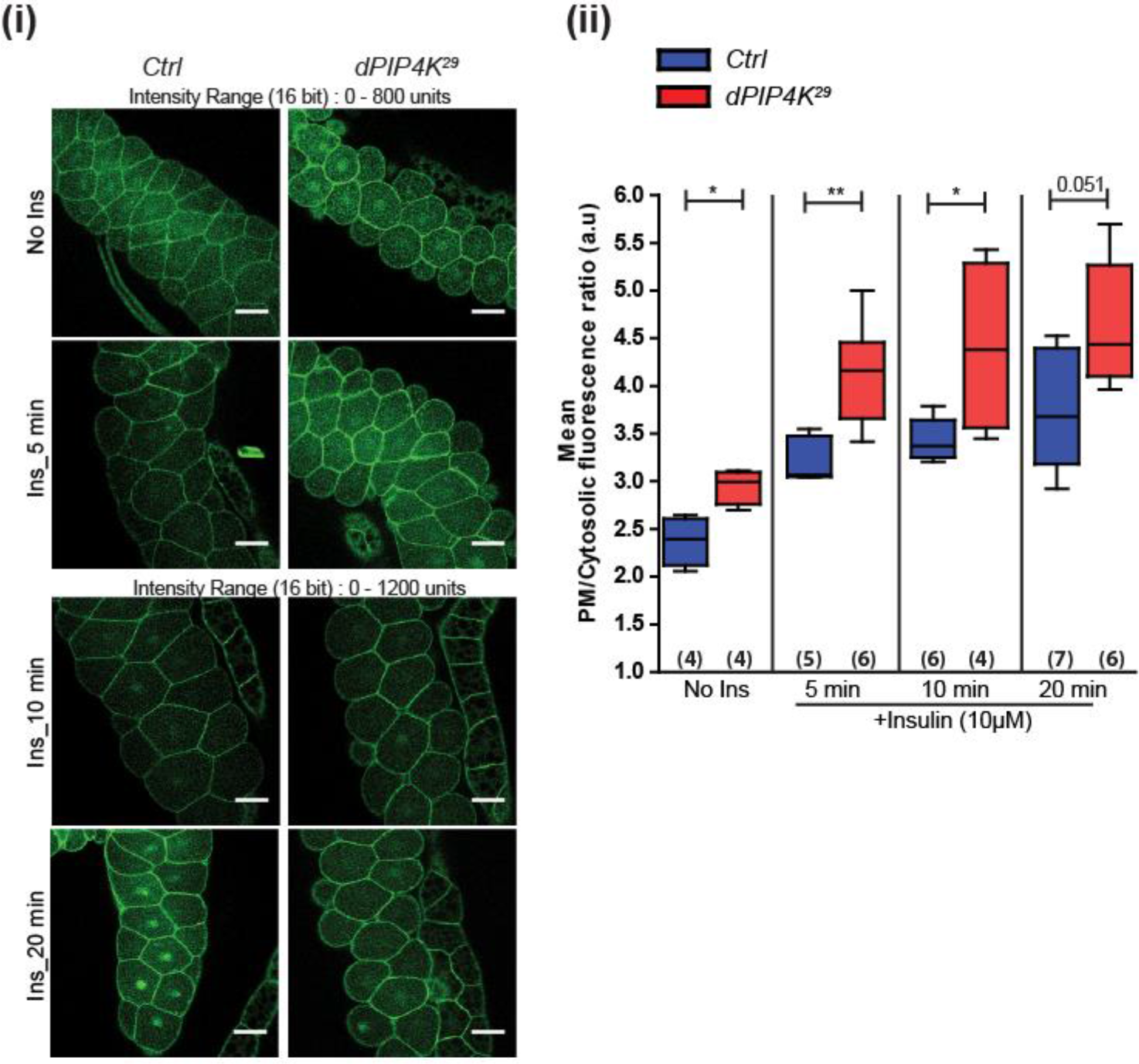
PIP_3_ measurement with increasing time of insulin stimulation. **A(i).** Confocal z-projections of salivary glands expressing GFP-PH-GRP1 in wildtype and *dPIP_4_K*^*29*^ backgrounds. Salivary glands dissected from wandering 3 rd instar larvae were stimulated or not with 10 μM bovine insulin for indicated times, fixed and imaged. **A(ii).** Relative PIP_3_ levels were measured as a ratio of mean fluorescence intensity at the plasma membrane to that in the cytosol. Whiskers in the box plots represent minimum and maximum values, with a line at the median. Numbers inside the parentheses below the plots indicate the no. of biological replicates used for the measurement. *Mann Whitney test* used for statistical analysis of the distributions. \*\**p-value* <0.01, \*\*\**p-value* < 0.001. Genotypes: *AB1Gal_4_, tGPH*/+ and *AB1Gal_4_, tGPH*/+; *dPIP_4_K*^*29*^

**Fig. S5.**
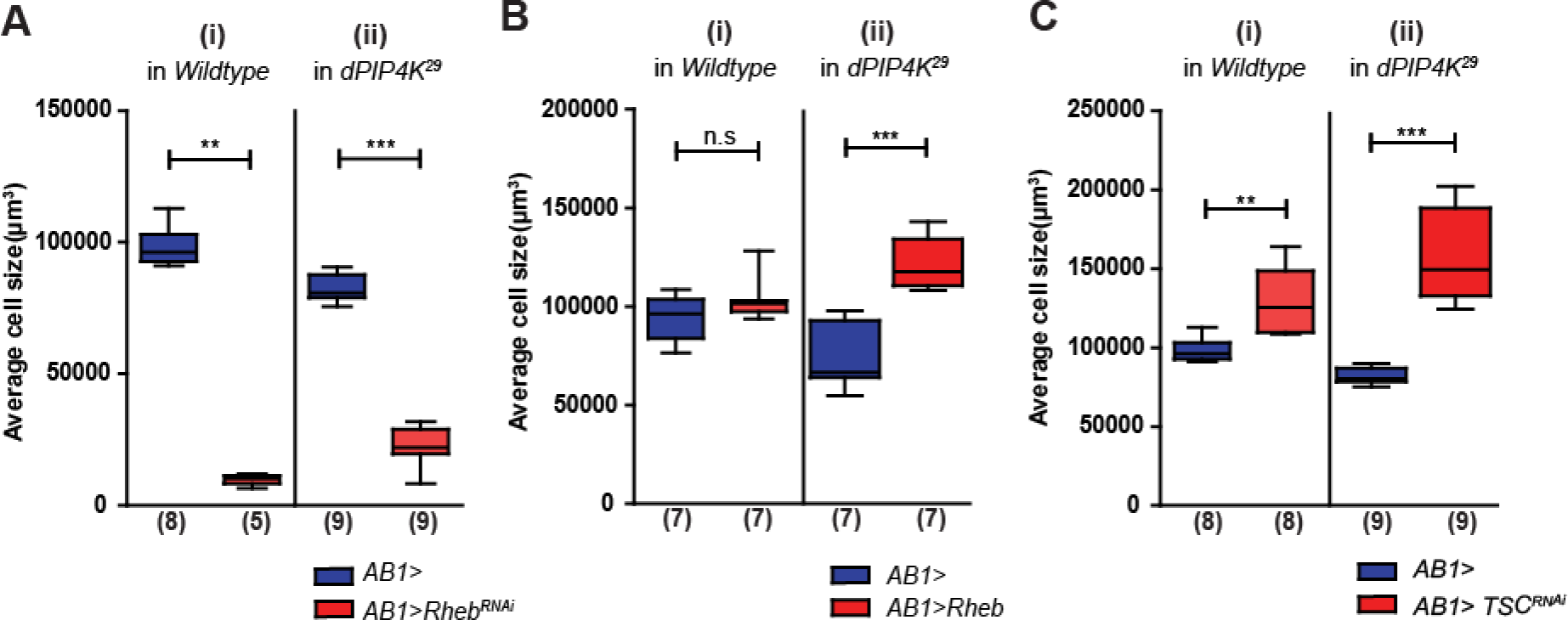
TORC_1_ activity regulates cell size in salivary glands. Cell size measurements in salivary glands upon **(A)** Knockdown of Rheb **(B)** overexpression of dRheb **(C)** Knockdown of TSC in *wildtype* and *dPIP_4_K*^*29*^ backgrounds. Whiskers in the box plots represent minimum and maximum values, with a line at the median. Numbers inside the parentheses below the plots indicate the no. of biological replicates used for the measurement. *Mann Whitney test* used for statistical analysis of the distributions. \*\**p-value* <0.01, \*\*\**p-value* <0.001. Genotypes: **A(i).** *AB1GC1I4*/+ *and AB1Gal_4_/UAS-Rheb*^*RNAi*^ and **(ii).** *AB1Gal_4_*/+; *dPIP_4_K*^*29*^ *and AB1Gal_4_/UAS-Rheb*^*RNAi*^; *dPIP_4_K*^*29*^ **B(i).** *AB1Gal_4_*/+ *and UAS-dRheb*/+; *AB1Gal_4_*/+ and **(ii).** *AB1Gal_4_*/+; *dPIP_4_K*^*29*^ *and UAS-dRheb*/+; *AB1Gal_4_*/+; *dPIP_4_K*^*29*^. **C(i).** *AB1Gal_4_*/+ *and UAS-Tsc1*^*RNAi*^/+; *AB1Gal_4_*/+ and **(ii).** *AB1Gal_4_*/+; *dPIP_4_K*^*29*^ *and UAS-Tsc1*^*RNAi*^/+; *AB1Gal_4_*/+; *dPIP_4_K*^*29*^. Controls for A(ii) and C(ii) represent values from the same dataset.

